# A Bioinert Hydrogel Framework for Precision 3D Cell Cultures: Advancing Automated High-Content and High-Throughput Drug Screening

**DOI:** 10.1101/2024.08.27.609940

**Authors:** Hyunsu Jeon, Tiago Thomaz Migliati Zanon, James Carpenter, Aliciana Ilias, Yamil Colón, Yichun Wang

## Abstract

Enhanced drug testing efficiency has driven the prominence of high-content (HC) and high-throughput (HT) screening (HCHTS) in drug discovery and development. However, traditional HCHTS in well-plates often lack complexity of *in vivo* conditions. 3D cell cultures, like cellular spheroids/organoids, offer a promising alternative by replicating *in vivo* conditions and improving the reliability of drug responses. Integrating spheroids/organoids into HCHTS requires strategies to ensure uniform formation, systemic function, and compatibility with analysis techniques. This study introduces an easy-to-fabricate, low-cost, safe, and scalable approach to create a bioinert hydrogel-based inverted colloidal crystal (BhiCC) framework for uniform and high-yield spheroid cultivation. Highly uniform alginate microgels were fabricated and assembled into a colloidal crystal template with controllable contact area, creating engineered void spaces and interconnecting channels within agarose-based BhiCC through the template degradation by alginate lyase and buffer. This results in a multi-layered iCC domain, enabling the generation of *in-vitro* 3D culture models with over 1,000 spheroids per well in a 96-well plate. The unique hexagonal-close-packed (HCP) geometry of iCC structure enables HCHTS through conventional plate reader analysis and fluorescent microscopy assisted by house-developed automated data processing algorithm. This advancement offers promising applications in tissue engineering, disease modeling, and drug development in biomedical research.

## 1. Introduction

Human disease models have become increasingly important in drug development due to the high failure rate of drug candidates in clinical trials. Despite a 44% increase in R&D expenditures by the top 15 pharma companies since 2016, reaching $133 billion in 2021, the drug attrition rate hit an all-time high of 95% that year.^[1]^ Most drugs fail in clinical stages despite proven efficacy and safety in animal models, largely because clinical trial decisions rely almost exclusively on animal-derived data. This translational gap has led to no significant increase in new drug approvals despite growing investments in pharmaceutical R&D.^[2]^ While human disease models based on 2D culture have widely been used for high-content (HC) and high-throughput (HT) screening (HCHTS), 2D culture environments offer cells a relatively simple and uniform environment.^[3,4]^ Such models are not sufficient to represent complex phenomena, such as tissue structures,^[5,6]^ spatial cell populations,^[7,8]^ and dynamic physiological processes,^[9,10]^ leading to differences in drug deliveries and efficacies.^[11,12]^ Hence, there has been a critical need to develop more reliable and accurate human disease models *in vitro* to better mimic the native environment, including tissue architecture in 3D coordinates, cell-cell, cell-extracellular matrix (ECM), and ECM-drug interactions.^[13]^

To address the translational gap, 3D cell culture techniques have emerged as promising solutions, with various human disease models developed over recent decades, including cellular spheroids, organoids, and scaffolded tissue models.^[3]^ Cellular spheroids and organoids, as scaffold-free 3D micro-physiological systems, effectively replicate tissue structures and functions, creating a more physiologically relevant environment for cellular activities.^[9]^ By spontaneously mimicking *in vivo* conditions through the expression of ECM, they not only promote physiologically reliable cellular behavior (*e.g*., cell differentiation, proliferation, signaling, and drug response in *in-vivo* conditions)^[14,15]^ but also allow dynamically precise molecular behavior (*e.g*., diffusion, transport, accumulation, and drug delivery within the 3D cellular system).^[12,16]^ Furthermore, the spherical geometry of cellular spheroids offers a straightforward 3D spatial coordinate system that closely mimics real tissue, representing cell-cell, cell-ECM, or ECM-ECM interactions.^[17]^ This simplification facilitates efficient monitoring/tracking and mathematical modeling of drug transport and efficacies as well as cell responses within the 3D cellular environment.^[18–20]^ To achieve 3D HCHTS that are compatible with traditional drug screening methods^[3,21]^ and adaptable to advanced artificial intelligence (AI) or machine learning (ML) analysis,^[3,22]^ various methodologies, such as hanging-drop method,^[23]^ bioreactor culture,^[24]^ suspension culture,^[25]^ and well-plate-based pellet culture,^[26]^ have been introduced to parallelize the generation, culturing, and analysis of spheroids and organoids. However, these methods exhibit spheroid throughput/content scalability limitations and/or spheroid size dispersity,^[27]^ showing a notable trade-off between the two features. Additionally, many of these methodologies for 3D HCHTS require complicated operation or are not compatible with the conventional readouts for drug testing, further complicating their adoption in both static and dynamic systemic analysis (*e.g*., endpoint or one-point analysis; *in-situ*, continuous, or real-time analysis).^[3,28]^

To reduce the complexity of workflow, platforms for *in-situ* spheroid generation and culture have been developed, which have aimed to provide specific geometric arrangements to spheroids in high-yield, with uniform and controlled size. Representatively, stemmed from the colloidal crystal topology,^[29–31]^ Lee *et. al*. introduced inverted colloidal crystal (iCC) structures as a high-yield and uniform spheroid culture platform,^[32–37]^ made of polyacrylamide (PAA) hydrogel backbone templated from polystyrene (PS) microbeads, offering uniform and hexagonal-close-packed (HCP) spherical void spaces interconnected with channels. The HCP geometry in the iCC hydrogel ensures interconnectivity among all spherical cavities and achieves a maximum theoretical porosity of 74%,^[32]^ enabling optimal packing of spheroids during their generation, maturation, and arrangement *in situ*. Inspired by this approach for *in-situ* formation of 3D *in vitro* models, various studies have suggested iCC framework-based tissue models using different combinations of template and backbone materials. Among various candidate template materials, including inorganic, organic, composite, and polymeric types,^[38,39]^ most studies have used microbeads made of polystyrene (PS), polymethyl methacrylate (PMMA), or glass due to their size ranges appropriate for cellular spheroid/organoid generation (*e.g*., 100–500 μm).^[36,40–42]^ However, these materials are commercially expensive, and require specialized instrumentation for large-scale fabrication.^[43]^ Moreover, their inelastic nature requires additional steps for channel interconnection after HCP assembly, such as precise high-temperature control (*e.g*., melting temperatures: >140 °C for PMMA, >240 °C for PS, and >700 °C for glass),^[29,40,44]^ which can introduce further hazards in framework fabrication. Furthermore, their chemically robust nature makes removal after backbone casting challenging and potentially hazardous, often requiring harsh degradation conditions and lengthy purification processes (*e.g*., tetrahydrofuran (THF), dichloromethane (DCM), hydrochloric acid (HCl), and/or hydrofluoric acid (HF)).^[45,46]^ Collectively, despite these advancements of iCC frameworks, the complexity and cost of the materials, along with the need for specialized procedures, present significant challenges for their widespread adoption.

Our work introduces a bioinert hydrogel-based iCC (BhiCC) framework that meets key requirements for 3D HCHTS, including uniform and high-yield cancer spheroid generation, facile testing, and automated analysis. Besides, it addresses the challenges of the iCC platform by enhancing scalability, cost-effectiveness, safety, and ease of fabrication. Specifically engineered framework fabrication process employs facile forming of HCP assembly of alginate microgels with controllable and spontaneous contact area through stiffness control of microgel, serving as the primary template for the iCC domain. The combinatorial degradation of microgels using enzyme and phosphate buffered saline (PBS) successfully creates void spaces within the agarose-based iCC without harsh or hazardous conditions, resulting in a BhiCC framework with multiple HCP layers and interconnected channels (**Figure 1A**). The framework demonstrates the consistent, uniform, and high-yield spheroid generation regardless of cancer types, as HC and physiologically reliable *in-vitro* models (*e.g*., >150 spheroids μL^-1^; > 1,000 spheroids per well in a 96-well plate; ECM expression; **Figure 1B**). Moreover, the unique HCP geometry of the iCC structure enables the integration of this framework with both conventional drug testing with a plate reader as a proof-of-concept HT drug testing (**Figure 1C**) and fluorescent microscopy-based automated image processing as a proof-of-concept HC drug testing (**Figure 1D**), demonstrating the successful implementation of a reliable 3D HCHTS platform.

**Figure 1.**
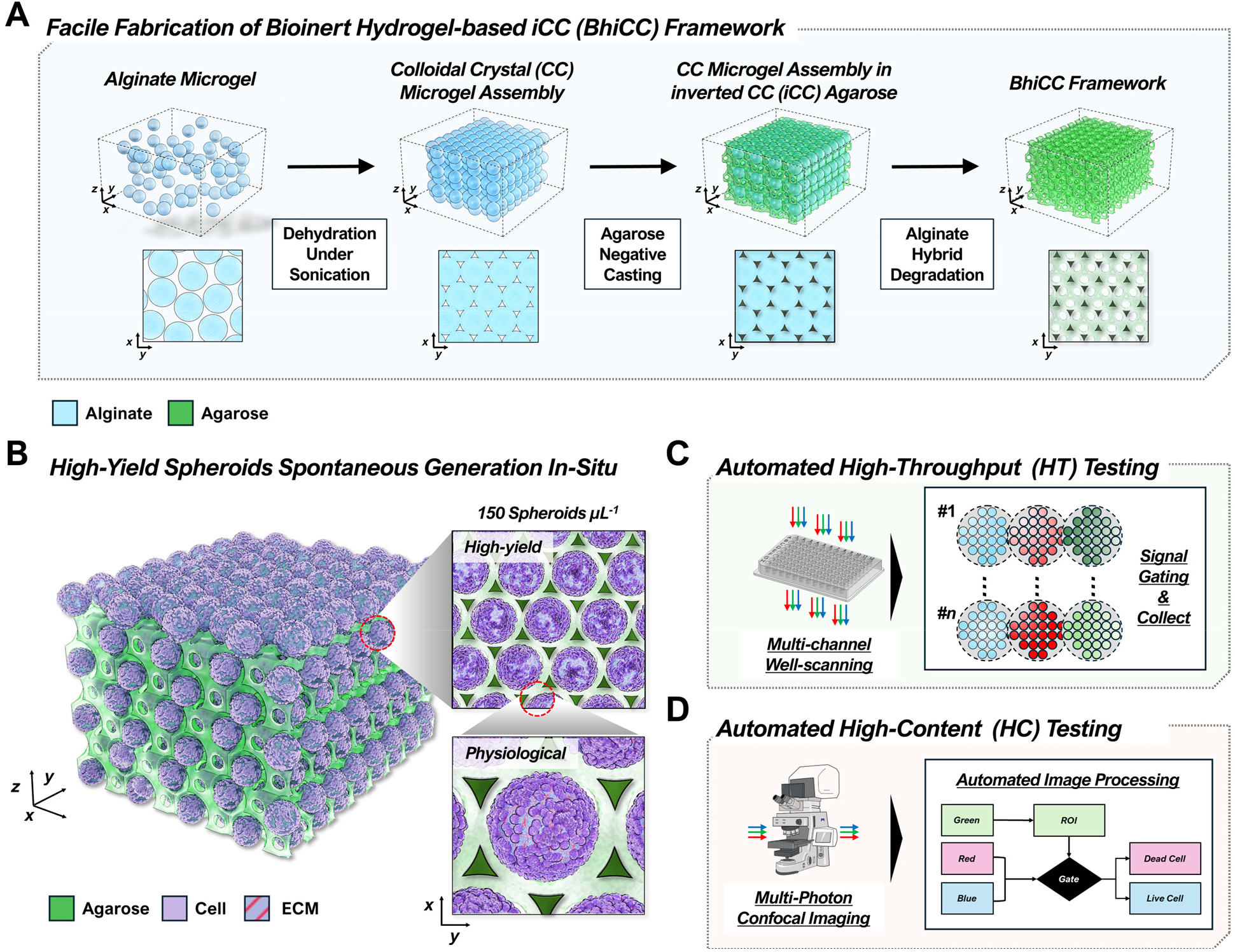
Schematic illustration of the workflow for high-throughput (HT) and high-content (HC) drug testing using the bioinert hydrogel-based inverted colloidal crystal (BhiCC) framework. **(A)** Schematic showing the BhiCC framework fabrication process. The concurrent use of alginate microgel and agarose hydrogel facilitated the safe, cost-effective, and straightforward generation of the BhiCC framework. **(B)** Spontaneous generation of highly uniform 3D spheroids within the BhiCC framework produced with high yield and demonstrating physiological mimicry of tumor tissue, including ECM expression. **(C-D)** Schematic flow outlining proof-of-concept **(C)** HT and **(D)** HC drug screening approaches using the BhiCC framework in HT and/or HC domains.

## 2. Result and Discussion

### 2.1 Fabrication and Mechanical Characterization of Alginate Microgels

Alginate hydrogel, made from alginic acid and calcium ions (Ca^2+^), was chosen as the materials of the primary template for the BhiCC framework due to not only its biodegradability, biocompatibility (*e.g*., both for itself and its degradates), and rapid gelation under mild conditions,^[47]^ but also tunable mechanical properties for HCP assembly and controllable contact areas of the primary template. To understand the mechanical behavior of alginate hydrogel, mechanical tests, including rheological analysis, were performed within a range of alginate contents in alginic acid solution (*e.g*., *C*_*alg*_; 2.0, 2.5, and 3.0%_w/v_). From the strain-sweep rheology tests (**Figure S1**), all the alginate hydrogels showed higher storage modulus (*i.e*., *G’*; 30–50 kPa) than loss modulus (*i.e*., *G”*; 5–10 kPa) under a shear strain range (*i.e*., *ε*_*S*_; 0.01–1%), demonstrating their gel-like mechanical behavior. The dynamic mechanical analysis (DMA) for a compression setting was performed on alginate hydrogels in three concentrations to understand further the mechanical behavior of alginate hydrogel, especially under compression. As a result, the compressive stress (*i.e*., *σ*_*C*_) *versus* compressive strain (*i.e*., *ε*_*C*_) curves for all alginate hydrogels were significantly dependent on *C*_*alg*_ on the hydrogel phases (**Figure 2A** and **Figure S2A**). Notably, the compressive modulus (*i.e*., *E*_*C*_), estimated from the slope at *ε*_*C*_ <0.1% following the relationship 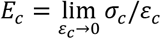, increased *C*_*alg*_ dependently (*e.g*., 2.0%_w/v_: 477.4±53.1 kPa; 2.5%_w/v_: 1128.3±13.7 kPa; 3.0%_w/v_: 2156.0±151.9 µm; **Figure 2B** and **Figure S2B**). This feature implies that the alginate hydrogel with lower *C*_*alg*_ behaves more elastically and deforms easily in a compressive direction, while higher *C*_*alg*_ shows less deformation.

**Figure 2.**
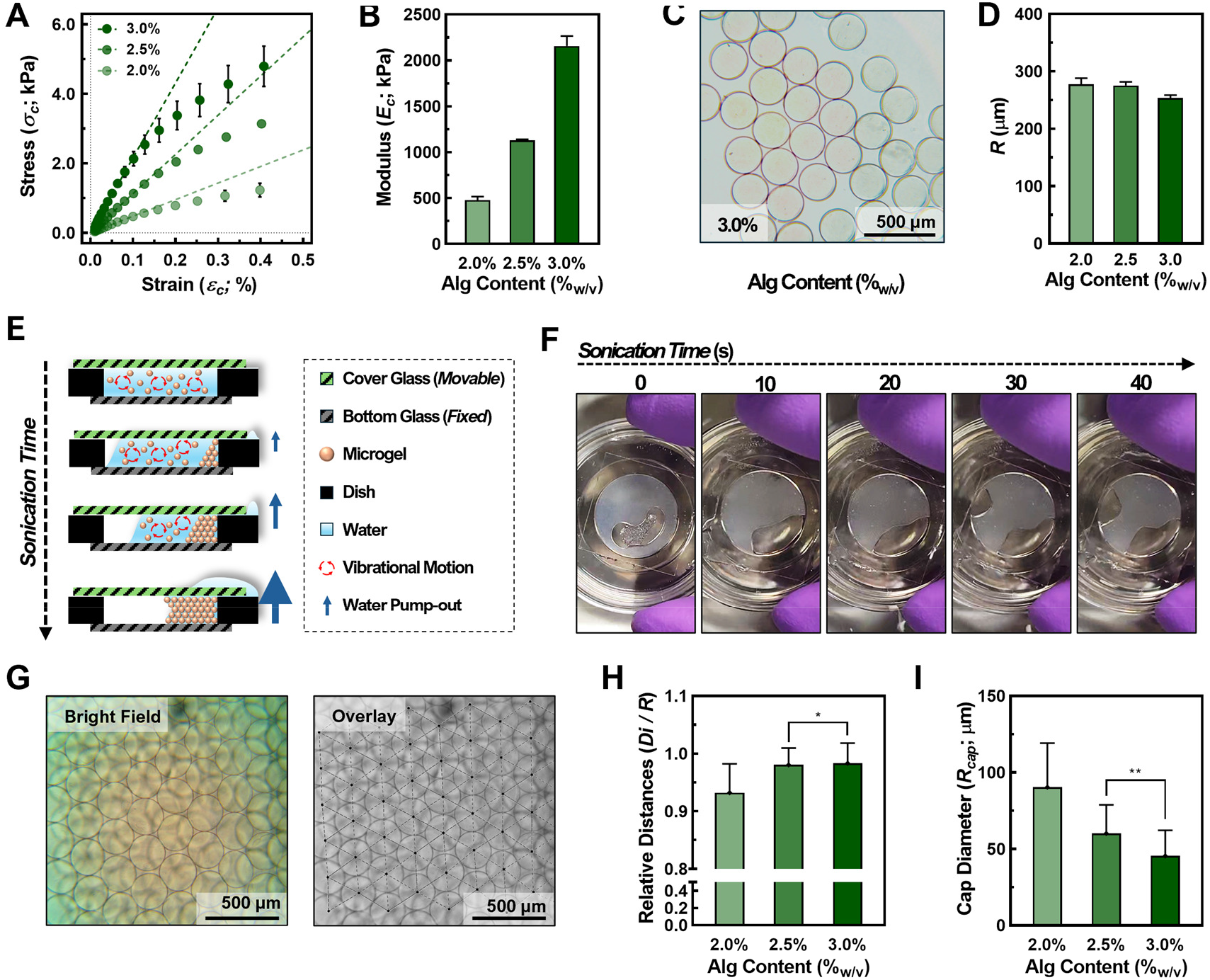
Hexagonal closed packed (HCP) formation with alginate microgel. **(A)** Compressive stress *σ*_*C*_ -strain *ε*_*C*_ curve for alginate hydrogels (*C*_*alg*_: 2.0, 2.5, and 3.0%_w/v_; error bar: ±SD; *N*=3). **(B)** Compressive modulus *E*_*C*_ of alginate hydrogels estimated from *σ*_*C*_-*ε*_*C*_ curves (error bar: ±SD; *N*=3). **(C)** Microscopic images of alginate microgel fabricated from electrospraying technique (*C*_*alg*_: 3.0%_w/v_; scale bar: 500 μm). **(D)** Diameter of alginate microgels (*C*_*alg*_: 2.0, 2.5, and 3.0%_w/v_). **(E)** Scheme illustrating dehydration of microgel in pseudo-slit system forming HCP assembly. **(F)** Time-lapse digital images showing HCP assembly formation. **(G)** Microscopic images of HCP assembly made of microgel assembly (left: bright-field image and right: overlayed image with HCP analysis; scale bar: 500 μm). **(H)** Relative distances between particles (*Di/R*; error bar: ±SD; *N*=60; \**P*<0.05 from unpaired t test). **(I)** Estimated spheroid cap diameters (*R*_*cap*_; error bar: ±SD; *N*=60;\*\**P*<0.01 from unpaired t test).

As structure units of HCP assembly as the primary template for the BhiCC framework, alginate microgels with *C*_*alg*_ ranges were prepared *via* the electrospraying technique (**Figure 2C** and **Figure S3A**). As a result, all microgel groups across *C*_*alg*_ ranges consistent size uniformity (*e.g*., 2.0%_w/v_: 277.2±10.8 µm; 2.5%_w/v_: 275.1±6.4 µm; 3.0%_w/v_: 253.3±5.1 µm; **Figure 2D**), with low standard deviations (SDs) (*e.g*., 3.89%, 2.33%, and 2.01%, respectively), qualified for HCP assembly (*e.g*., 1–5% in previous reports^[40]^). The aspect ratios for all microgel groups showed close-to-1 values (*e.g*., 1.059±0.049, 1.048±0.038, 1.023±0.022, respectively; **Figure S3B**), demonstrating the high roundness of microgels as spheres. Consequently, these results showed excellent uniformity and roundness of alginate microgels, which enables the formation of HCP geometry under close-packed assembly as the primary template for the desired BhiCC framework.

### 2.2 Assembling Alginate Microgels into Primary HCP Microgel Template

To achieve the HCP assembly of microgels for the BhiCC framework, we developed a simple approach: dehydration under a pseudo-isolated slit (*e.g*., **s**ee *Experimental Section*; **Figure 2E** and **Figure S4A**). Herein, 1.5×10^4^ microgels were positioned at the bottom of a 15 mm glass-bottom dish (*e.g*., Inner diameter: 15 mm; Outer diameter: 35 mm) and gently covered with the cover glass, forming a pseudo-isolated slit. Under the sonication-inducing vibrational motion of microgels, the cover glass confined microgels within the slit while still permitting water to escape, allowing the vibrations to prompt dehydration inside the slit, which restricted the microgel motion while compelling them into closed packing. Consequently, within 1 min of the sonication process, the microgels were successfully separated from the liquid phase, left inside the slit, while water embedding microgels was pumped outside of the slit (**Figure 2F**). This dehydration process spontaneously resulted in the closest packing of microgels as the HCP microgel template (**Figure 2G** and **Figure S4B**).

Notably, we innovate the channel formation procedure from that of the previous works to significantly improve the procedural simplicity and safety (*e.g*., no need for high temperature, toxic solvent, or secondary coating ^[40,43,48]^), leveraging the compression-responsive feature of the microgels that induced symmetric deformation. Such features lead to the close contacts of sphere caps and their deformation perpendicular to the contacting planes between microgels within the HCP microgel template, which is the key factor for the controllability in channel diameter within the BhiCC framework. This microgel cap contact-induced deformation was influenced by the force balance between gravitational force (*i.e*., *F*_*g*_), buoyant force *F*_*b*_, and compressive stress-mediated forces *F*_*C*_ (*e.g*., **Equation (1)**), which led to the relationship described in **Equation (2)** and **Equation (3)** (*e.g*., where *R*: Microgel diameter, *Di* : Distance between microgels, *p*_*w*_ : Water density, and *g* : Gravitational acceleration; see **Supplementary Note 1**). This relationship indicates the straight correspondence between *E*_*C*_ and spontaneously formed sphere cap sizes (*R*_*Cap*_) within the microgel assembly (*i.e*., interconnecting channel diameters in BhiCC framework) (*e.g*., see **Equation (1)**–**Equation (4)**).

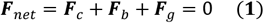

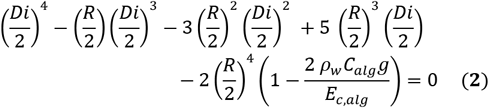

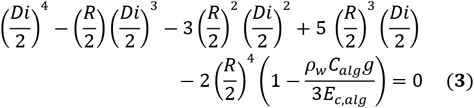

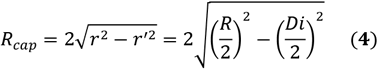

As a proof of estimation, the center-to-center distances between the microgels within the microgel assembly were recognized and collected through image processing (**Figure S4B**; See **Figure 2G**) and represented as relative distances (*e.g*., *Di/R*; **Figure 2H**) for *C*_*alg*_ respectively. The smallest relative distance value was observed in the microgel assembly with a lower *C*_*alg*_while they increased to 1 with a higher *C*_*alg*_ (*e.g*., 0.932±0.050, 0.980±0.029, 0.983±0.034, respectively) with statistical significance. Consequently, *R*_*Cap*_, estimated from the relationship 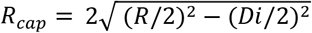, showed the opposite *C*_*alg*_ correspondence (*e.g*., 90.3±28.8 µm, 60.1±18.7 µm, 45.4±16.7 µm, respectively; **Figure 2I**), implying the controllability in channel diameter within the BhiCC framework. Overall, the newly developed approach for HCP assembly of microgels by dehydration under the pseudo-isolated slit are easy, fast, and effective, while the controllable and uniform contact areas between the microgels in this HCP template can be tuned by the mechanical properties of the hydrogel materials to engineer interconnected channels in the final BhiCC frameworks.

While various methods have been reported for the self-assembly of microspheres (*i.e*., microparticles or microbeads) in an HCP domain, our approach offers a faster, simpler, more cost-effective, and safer alternative. Previous methods ^[40,43,48]^ including the ones from our previous reports ^[30,42,49–51]^ typically involve inelastic microspheres made from inorganic, organic, composite, or polymeric materials with high monodispersity. These methods often require toxic and hazardous synthesis conditions in their fabrication (*e.g*., reduction, hydrolysis, or hydrothermal reactions), which demand toxic organic solvents, harsh pH, and/or high temperatures.^[40]^ Additionally, due to the inelastic nature of these microspheres, high-temperature treatment (*e.g*., *T*_*m*_ ^[29,40,44]^) are often necessary to establish interconnected contact area between microspheres. In contrast, our approach utilizing elastic alginate microgels involves a straightforward microgel fabrication process involving only alginic acid, and Ca^2*+*^ ions under mild conditions (*e.g*., well-established, simple, and orthogonal reaction under room temperature, aqueous solution, and mild pH ^[52]^). Moreover, our method not only provides tunable mechanical properties but also enables spontaneous interconnectivity of the microspheres through gravity alone due to their elastic nature.

On the other hand, various approaches have been introduced to assemble microbeads or microgels into HCP assembly, such as solvent evaporation,^[45,53]^ application of external forces,^[54]^ or microfluidic techniques.^[55]^ These methods often require an extensive time range (*e.g*., hour-to-day scale), specialized equipment (*e.g*., spin-coaters, microfluidic devices, or electric field generators), complex microsphere functionalities (*e.g*., paramagnetic properties^[54]^), or have limited scalability in shape of resulting assembly (*e.g*., confined to vertical dimension^[40]^). In contrast, our method uses a pseudo-isolated slit for HCP assembly of alginate microgels requiring only a simple sonication device and a glass-bottom dish, achieving cylindrical packing of HCP assembly of > 30,000 alginate microgels in 5 planar layers in <1 min, without help of additional microgel functionalization.

### 2.3 High-Strength Low-Melting Agarose (LMA) Hydrogel as Secondary Backbone Material

LMA hydrogel was selected as a secondary backbone material for the resulting BhiCC framework, featuring its biocompatibility and ultra-low affinity to cell and ECM. The low affinity of LMA allows cells aggregate with each other without adhering to the surface of microcavities, which is essential for spheroid/organoid formation due to the mechanical and physical confinement, as well as to maintain long-term spheroid/organoid culture.^[56]^ To obtain a robust hydrogel structure for the BhiCC framework, LMA hydrogel was fabricated from the hydrated LMA powder, condensed into the LMA paste, followed by a mild heating/cooling process (*e.g*., ∼ 15.6%_w/v_; 70 °C 30 min and room temperature 30 min; see *Experimental Section*; **Figure S5A**). As a result, LMA hydrogel stably exhibited gel-like behavior from the rheology test (**Figure 3A**), with higher *G’* (*i.e*., ∼300 kPa) than *G”* (*i.e*., ∼10 kPa) under a *ε*_*S*_ range (*i.e*., 0.01–1%). The observed *G’* value of approximately 300 kPa demonstrates that incorporating LMA not only offers great physical confinement for spheroid generation and enables higher mechanical robustness in manufacturing BhiCC framework, compared to other common backbone hydrogels for iCC structure previously reported,^[40,43,48]^ but also suggests its potential to replicate the mechanical properties of various tissue types near tumors (*e.g*., arterial walls, or veins^[57]^). To track its spatiotemporal information, we tagged fluorescein to the LMA (*i.e*., F-LMA; see *Experimental Section*) and observed materials under a fluorescence microscope. A clear fluorescence signal can be observed from the tagged fluorescein (*e.g*., Excitation: 492 nm and emission: 516 nm^[58]^) with hydrogel samples from only 10% tagging (*e.g*., 90% LMA: 10% F-LMA; **Figure 3B**). Moreover, during the temperature sweep conducted using a rheometer (**Figure 3C**), LMA and F-LMA hydrogel exhibited a loss of viscoelasticity at approximately 48 °C (*e.g*., *T*_*m*_; **Figure 3C**), indicating its retention of thermoplastic property, and thus moldability (**Figure S5B** and **Figure S6A**). Notably, the LMA hydrogel synthesized through our approach demonstrated superior mechanical robustness compared to other hydrogels previously reported as backbones for iCC structures^[41,42,59–63]^ or as agarose-based 3D cell culture platforms.^[64–66]^ This property allows the BhiCC framework to efficiently provide physical confinement for spheroid generation, while also ensuring robust scalability and stable integration with conventional assays, making it suitable for designing next-generation 3D HCHT drug screening platforms.

**Figure 3.**
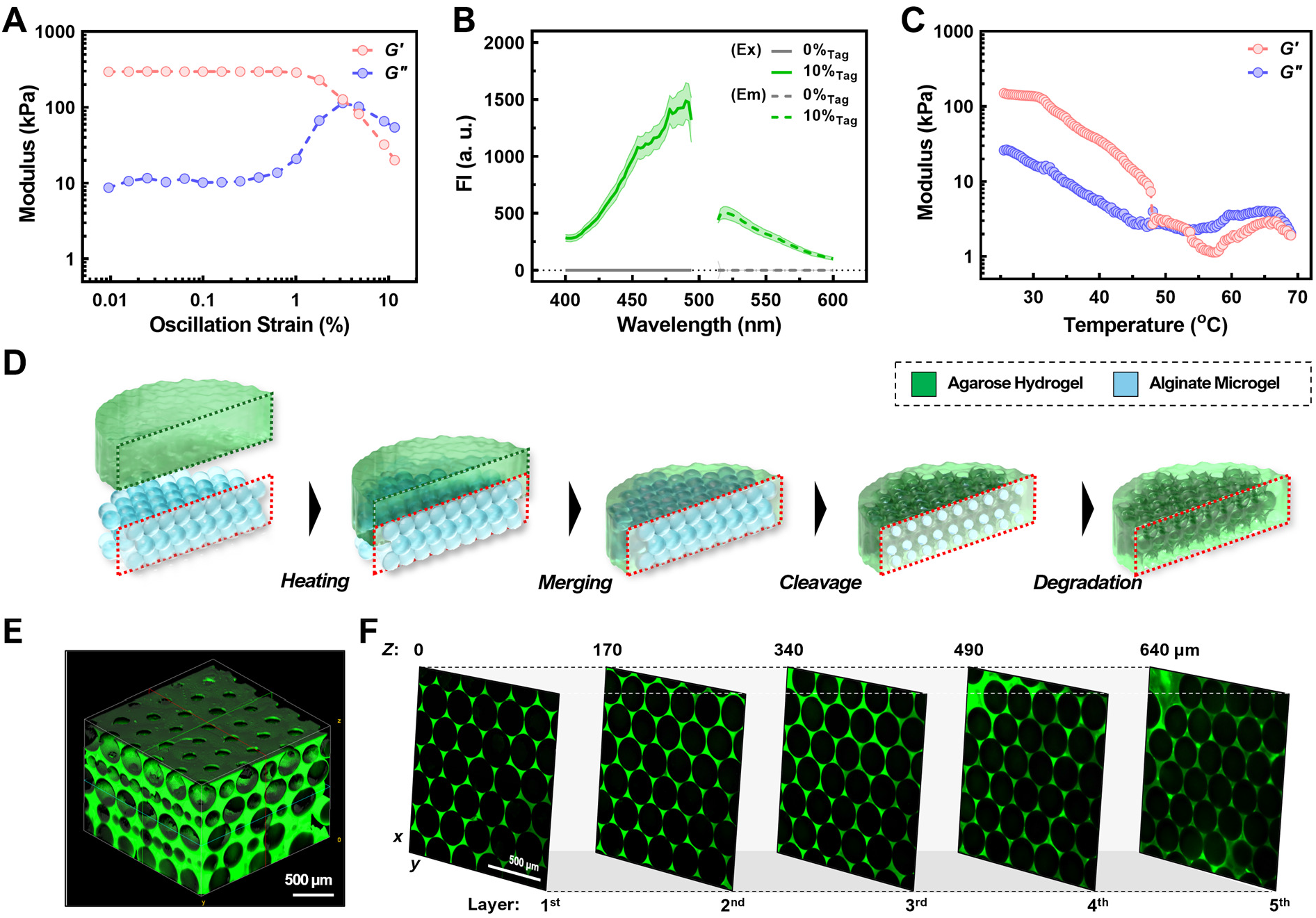
Fabrication of Bioinert hydrogel-based inverted colloidal crystal (BhiCC) framework from alginate microgel HCP template and low-melting agarose (LMA) primary backbone hydrogel. **(A)** Storage (*G’*; red curve) and loss (*G”*; blue curve) modulus of LMA hydrogel (15.6%_w/v_) under strain sweep at a fixed *f* of 1 rad/s. **(B)** Fluorescence measurement from 5-DTAF-tagged LMA (F-LMA) hydrogel (excitation: 492 nm and emission: 516 nm; error bar: ±SD; *N*=3). **(C)** Thermo-responsiveness of LMA hydrogel (*G’* as red curve and *G”* as blue curve; 1% *E*_*S*_; 1 rad/s *f*). **(D)** Schematic workflow to fabricate BhiCC framework. (E) 3D constructed two-photon confocal microscopic (TP-CLSM) image of resulting BhiCC framework (scale bar: 500 μm; green: 5-DTAF from F-LMA). **(F)** Multi-layered HCP formation within BhiCC framework demonstrated with single *Z*-plane TP-CLSM images (scale bar: 500 μm; green: 5-DTAF from F-LMA).

### 2.4 High-Strength Low-Melting Agarose (LMA) Hydrogel as Secondary Backbone Material

Given all features of alginate microgel template and LMA hydrogel, we designed facile methodology to fabricate BhiCC framework (**Figure 3D)**. In brief, LMA hydrogel was first positioned onto the HCP template assembled by alginate microgels and then thermally led the two phases merging at *T*=70 °C (*i.e*., *T*>*T*_*m*_; See **Figure 3C**), followed by solidification at room temperature. The LMA hydrogel, pre-shaped to the glass-bottom dish (**Figure S6B**), kept the HCP assembly stable during its infiltration, enabling successful HCP assembly-embedding hydrogel formation (**Figure S6C**), except in the 2.0%_w/v_ case due to poor LMA infiltration. Notably, the LMA encapsulated 5 layers of the HCP template, spontaneously exposing the sphere caps of 1^st^ layer outside the LMA hydrogel, which will generate top holes after microgel degradation, enabling cell infiltration into the final BhiCC hydrogel. To form the final BhiCC framework, the alginate microgel HCP template was first degraded enzymatically with alginate lyase (**(i)**–**(ii)** in **Figure S7A**), followed by chemical chelation of Ca^2+^ using PBS (**(ii)**–**(iii)** in **Figure S7A**). Consequently, microgel shrinkage was observed after cleavage (**(ii)** in **Figure S7B**), indicating successful separation of HCP assembly from the LMA hydrogel without deforming. After Ca^2+^ chelation, the microgel residue swelled gradually and was eliminated eventually, leaving the resulting BhiCC framework (**(iii)** in **Figure S7B**).

To verify the successful fabrication of uniform void spaces and interstitial channels within the BhiCC framework, we utilized two-photon confocal laser scanning microscopy (TP-CLSM) to track LMA phase in the BhiCC framework. As a result, the BhiCC framework exhibited a clear array of HCP crystalline void spaces (*e.g*., ∼250 µm in diameter) with uniform interconnecting channels between them **(Figure 3E** and **Figure S8A)**. Based on our theoretical estimation using a circle-in-circle close packing model, the expected number of void spaces in a single layer of the framework ranges from 3145 to 3155, which shows the high feasibility of our framework for 3D HC drug screening (*e.g*., see **Supplementary Note 2**). The channel diameters were 44.05±5.44 µm at the top side of the framework (*e.g*., the most bottom layer in fabrication process) while the side channels were 39.99±7.07 µm (**Figure S8B**). The diameters of both channels were sufficient to accommodate not just individual cells (*e.g*., 10–20 µm^[67]^) infiltrating the framework, but also to facilitate the exchange of nutrients and waste (*e.g*., sub-nanometric scale) for formation of spheroids or organoids. Moreover, the framework contained numerous layers of void spaces and channels within the HCP domain (*e.g*., 5 layers; **Figure 3F**) and exhibited consistent and periodic fluorescence intensity (FI) patterns along the *Z* position (**Figure S8C**).

### 2.5 Spontaneous Generation of Cellular Spheroids in a High-Yield Domain

To generate high-yield spheroid structures in the BhiCC framework, we develop a centrifugal method for cell seeding. As illustrated in **Figure 4A**, a customized polydimethylsiloxane (PDMS) holder, integrated within a commercial 50 mL centrifuge tube, was used to facilitate perpendicular cell penetration through the top-side channels of the BhiCC framework. For the proof-of-concept, we first employ human hepatoma cells (*i.e*., HepG2) to generate cellular spheroids. The serial microscopic images of a HepG2-seeded framework showed the process of the cell condensation into spheroids (**Figure 4B**). The seeded cells were initially observed as individual suspension cells or loose cellular aggregates (*e.g*., see 0 h in **Figure 4B**), while started showing smooth surface without distinguishable cells and formed solid spherical shapes after 72 h as cellular spheroids. This spontaneous and successful spheroid formation was further demonstrated after 144 h from the cell seeding with a multi-photon microscopy while utilizing 5-DTAF-tagged framework (*e.g*., containing F-LMA) (*e.g*., green channel) and CMTPX as a live cell tracker (*e.g*., red channel) (**Figure 4C**). We further expanded the applications to other common cancer cell lines (**Figure 4D, Figure S9**, and **Figure S10**): human prostate carcinoma (*i.e*., LNCaP), human adenocarcinoma (*i.e*., MCF-7), and human neuroblastoma (*i.e*., SH-SY5Y). All of these cell lines can form condensed cellular spheroids gradually (**Figure S9**), with high yield after 120 h from cell seeding (**Figure 4D** and **Figure S10**). These results demonstrated that our method is capable of highly uniform spheroid generation in high-yield spontaneously, and most importantly it is cell-type independent that can apply to 3D HCHTS for diverse diseases.

**Figure 4.**
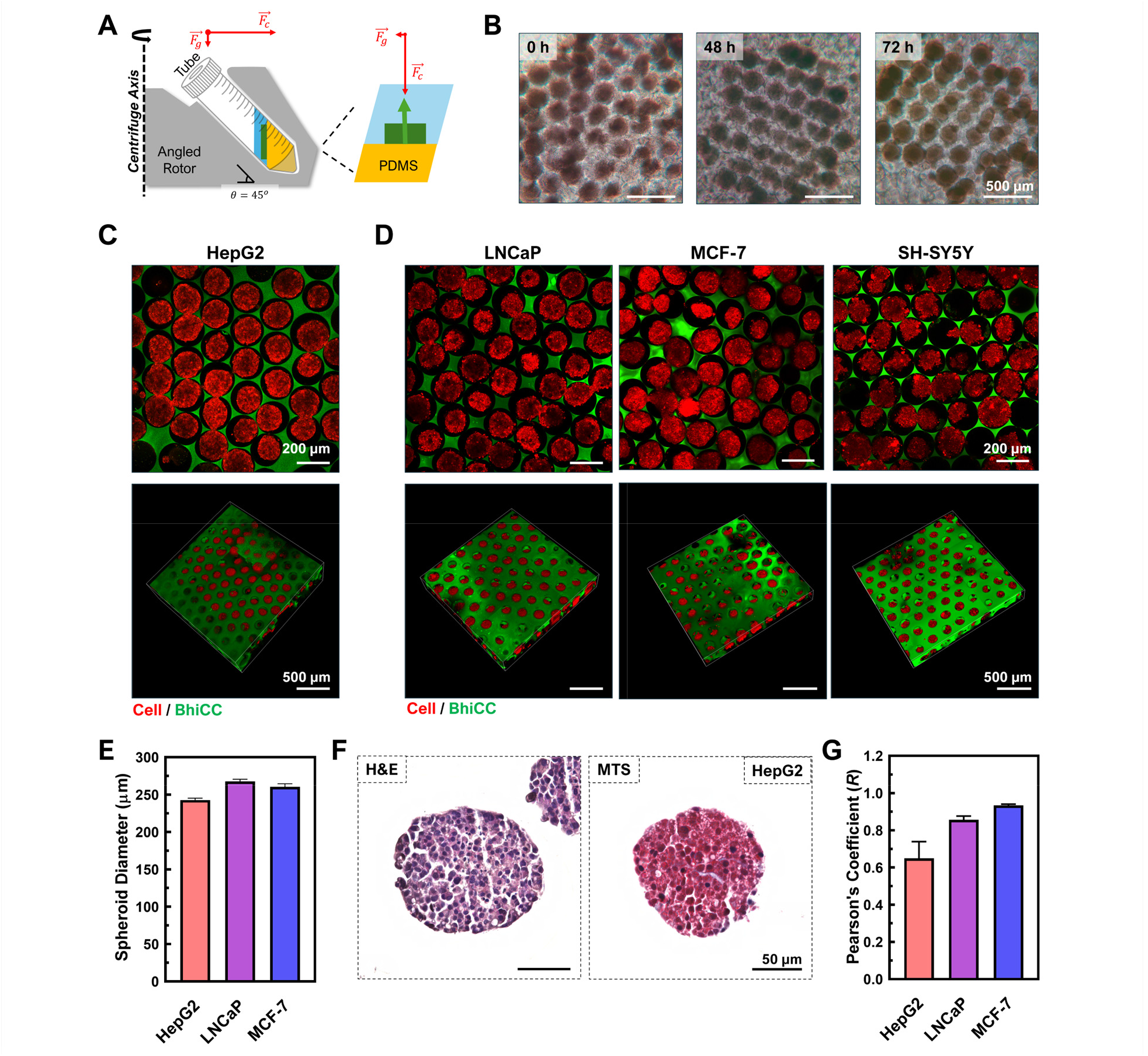
High-yield spheroid generation in BhiCC framework. **(A)** Scheme describing centrifugal approach for cell seeding in BhiCC framework. **(B)** Time-lapse microscopic images of HepG2 spheroid generation in BhiCC framework (24, 48, and 72 h; scale bar: 500 μm). **(C)** TP-CLSM images of HepG2 spheroids in BhiCC framework (top: single *Z*-plane image and bottom: 3D reconstructed image; scale bar: 200 μm (top) and 500 μm (bottom); green: 5-DTAF from F-LMA and red: CMTPX from live cells). **(D)** TP-CLSM images of LNCaP (left), MCF-7 (center), and SH-SY5Y (right) spheroids in BhiCC framework (top: single *Z*-plane image and bottom: 3D reconstructed image; scale bar: 200 μm (top) and 500 μm (bottom); green: 5-DTAF from F-LMA and red: CMTPX from live cells). **(E)** Spheroid diameters collected from HepG2, LNCaP, and MCF-7 TP-CLSM images (error bar: ±SD; *N*=100). **(F)** Histological result from HepG2 spheroids generated in BhiCC framework (left: H&E and right: MTS; 50 μm). **(G)** Comparison of Pearson’s coefficients for HepG2, LNCaP, and MCF-7 spheroids, derived from the two-channel correlation analysis of blue and red colors in MTS (error bars represent ±SD; *N*=4).

We quantify the sizes of spheroids by collecting the live cell signal (*i.e*., red channel) from TP-CLSM images. The diameter of the maximum area for each spheroid throughout *Z* depths (**Figure S11**) was selected to represent the diameter of spheroids (**Figure 4E**). Spheroids for each cell line showed specific diameters despite the same seeding density respectively due to their nature of growth (*e.g*., HepG2: 242.7±13.21 µm, LNCaP: 267.7±19.42 µm, MCF-7: 260.5±25.16 µm), while notably showing narrow size distributions for all cell lines (*e.g*., < 10% in SD for all cell lines). This fact further confirmed that our system can support high-yield uniform *in-vitro* model generation across various cancer types in high-yield and HC domain.

To characterize the structure and composition of cellular spheroids inside the BhiCC framework, we performed histological examinations. Here, we selected two approaches for staining spheroids: hematoxylin and eosin staining (*i.e*., H&E) and Masson’s Trichrome staining (*i.e*., MTS). The histological section from HepG2 spheroids (**Figure 4F** and **Figure S12A**) showed densely packed cellular structure under H&E (*e.g*., nuclei as deep-blue-purple; cytoplasm and ECM as orange-pink-red color^[68]^) while ECM structure connecting cellular structures can clearly be observed under MTS (*e.g*., ECM as blue; nuclei as dark red-purple; cytoplasm as red/pink^[69]^). Interestingly, when performed the same staining procedure to LNCaP (**Figure S12B**) and MCF-7 (**Figure S12C**), there were similar packing densities of cellular structures under H&E but different tendencies of ECM among these cell lines under MTS (*e.g*., blue). To quantify the amount of the ECM formed inside spheroids, we performed color deconvolution for each cellular spheroids and analyzed the correlation between red and blue channels as Pearson’s correlation coefficient (***R***; **Figure 4G**) and the two-channel histogram mapping (**Figure S13**). The correlation analysis of HepG2 spheroids showed smallest ***R*** while MCF-7 showed highest (*e.g*., HepG2: 0.649±0.090; LNCaP: 0.856±0.020; MCF-7: 0.933±0.007). Notably, ***R*** between two data set represents the correlation degree from 0 (*e.g*., no correlation) to 1 (*e.g*., identical), implying the most abundant ECM generation in case of HepG2, while MCF-7 has a minimum blue color among three cell lines. This tendency matched with the two-channel histogram plots (**Figure S13**), which shows the most blue-shifted scatters for pixels for HepG2 spheroids (*i.e*., left panel) while almost linear-like scatters from MCF-7 (*i.e*., right panel). To note, previous studies have highlighted that spheroid and organoid culture outcomes are highly dependent on the culture method—whether scaffold-based or scaffold-free^[28,70–72]^—and the origin and phenotype of the cell lines used.^[73–75]^ The BhiCC framework offers the advantage of applicability across various cell types and enhances the maturation of ECM expression in high-yield 3D tissue models. Consequently, we successfully confirmed that the cellular spheroids derived from the BhiCC framework could form densely packed structures that express ECM components, which mimics the *in-vivo* tumor tissues.

### 2.6 Proof-of-Concept 3D HT Drug Screening

Leveraging the high-yield spheroid generation capability of the BhiCC framework, we developed a proof-of-concept HT drug screening system utilizing common drug screening instrument such as a plate reader. Notably, the mechanical properties of BhiCC framework enabled customized fitting into a conventional microwell plate (**Figure 5A**). To obtain the information of drug efficacy on spheroids, we pre-stained HepG2 and LNCaP cell lines with a live cell tracker (*e.g*., blue fluorescence) before seeding them in the BhiCC framework (*e.g*., containing F-LMA; green fluorescence), followed by staining dead cells with BOBO-3 Iodide after drug treatment (*e.g*., red fluorescence; see *Experimental Section*). Doxorubicin (DOX) was used as a representative drug to treat the tumor spheroids (*e.g*., 0, 10, 20, 50, 100, and 200 μM, respectively) in this study. We captured all fluorescence signals from the BhiCC framework containing both live and dead cells (*e.g*., *N*=4 for each concentration and cell line) by fluorescence measurement from a plate reader. The blue signal showed insignificant changes across all experimental replicate groups as DOX concentration increased (**Figure S14A**). In contrast, the red signal gradually increased in intensity with higher DOX concentrations, reflected in the well-by-well averaged values shown in **Figure S14B**. Collectively, these results indicate the feasibility of the BhiCC framework serving as HT drug screening only with conventional reagents for fluorescence staining assays and a conventional microwell plate, showing its high compatibility with traditional drug screening. Moreover, to further increase the statistical reliability and predictability on drug responses, we employed the fluorescent well-scan mapping function in the plate reader (**Figure 5B**), increasing the number of readings per well and collecting spatial information (**Figure S15** and **Figure 5C**). This resulted in 36 data points per framework, providing 36 statistical replicates per sample. This approach allowed us to represent the frequency distributions for each point and enabled the fitting of Gaussian plots (**Figure S16**). Similar to the single-readout results (*e.g*., see **Figure S15**), the plots from the blue signal collection (*FI*_*blue*_) showed insignificant decreases with increasing DOX concentrations (**Figure S16A**). In contrast, the red signal plots (*FI*_*red*_) gradually shifted with increasing DOX concentrations (**Figure S16B**). Additionally, given that each blue and red signal embeds spatial information from the BhiCC framework, we represented the ratio of the dead cell signal to the live cell signal followed by a voxel-to-pixel correction (*i.e*., Corrected *FI*_*red*_*/FI*_*blue*_; see *Experimental Section*) (**Figure 5D**). A clear signal ratio shift from the BhiCC framework for both cell lines was observed depending on the administered DOX concentration. The extracted mean and SD for each Gaussian plot allowed us to estimate the median effective dose (*IC*_*50*_) for each cell line (*e.g*., *IC*_*50*_=25.69 μM for HepG2; *IC*_*50*_=20.45 μM for LNCaP; **Figure 5E**). HepG2 spheroids showed a higher *IC*_*50*_ than LNCaP spheroids while exhibiting different positive cooperativity (*e.g*., Hill Coefficient; *n*=1.310 for HepG2; *n*=0.9572 for LNCaP). These results imply increased statistical reliability than single-point readouts (*e.g*., see **Figure S14**) in extracting drug response data from our BhiCC framework, representing statistically meaningful response curves in a HT domain, with reliable coefficient extractions including *IC*_*50*_ and the Hill coefficient. Nonetheless, this plate reader-based approach cannot be considered an ideal platform that fully leverages the potency of the BhiCC framework: the readouts per well in the 96-well plate were limited to 27, while the BhiCC framework contained over 3,000 spheroids per well.

**Figure 5.**
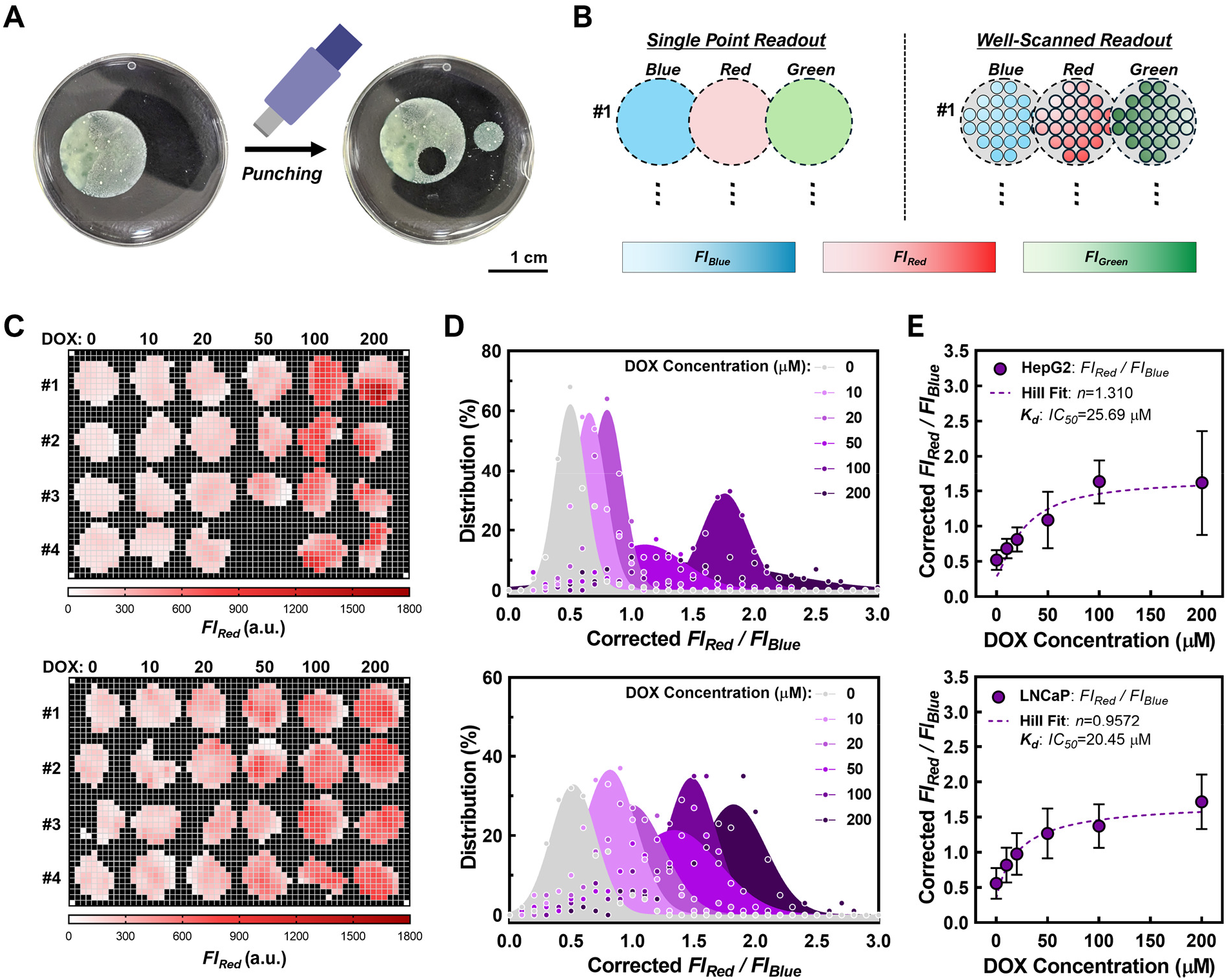
Proof-of-concept HT drug screening with BhiCC framework. **(A)** Facile cutting and fitting of BhiCC framework into a 96-well plate. **(B)** Scheme describing a well-scanning function of a microplate reader and working procedure for the BhiCC framework-based HT drug testing. **(C)** Heatmap of drug response result of live cell fluorescence intensity (*FI*_*blue*_) from BhiCC framework-embedded spheroids from HepG2 (top) and LNCaP (bottom) after DOX treatment (24 h treatment and 24 h incubation). Each column represents DOX concentrations from 0 to 200 µM, respectively, while first to fourth rows represent experimental replicates (*i.e*., Sample #1–#4; *N*=4). **(D)** Frequency histogram and Gaussian fitting of drug response results (referred as Corrected *FI*_*red*_*/FI*_*blue*_) of BhiCC framework-embedded spheroids derived from HepG2 (top) and LNCaP (bottom). **(E)** Drug response curves generated by average and SD values from each Gaussian fitting (top: HepG2 and bottom: LNCaP; error bar: ±SD; *N*=144).

### 2.7 Proof-of-Concept for 3D HC Drug Screening

To efficiently and effectively collect drug response data by leveraging 3D information of spheroids and the transparency of BhiCC framework with higher statistical reliability than the proof-of concept HT screening, we further utilized TP-CLSM to extract spatial information of each spheroid in the framework. We prepared DOX-treated HepG2 and LNCaP spheroids within the BhiCC framework, similar to the HT drug testing described in **Section 2.6** and performed TP-CLSM imaging with 4 imaging replicates (LNCaP: **Figure 6A** and HepG2: **Figure S17A**). Fluorescence signals from both blue live cells and red dead cells were successfully collected within the void spaces in the green BhiCC framework. Notably, every single image from the framework incorporated 50–80 HCP-arranged void spaces, each filled with cellular spheroids representing individual biological replicate. From the TP-CLSM images, LNCaP spheroids showed gradual dominating fluorescence signal changes from blue (*e.g*., live cells; high cell viability) to red (*e.g*., dead cells; low cell viability) with increasing DOX concentrations, while signals from HepG2 spheroids shifted more abruptly, especially in the range of 20–50 μM. These results imply that there were different drug responses between LNCaP and HepG2 spheroids to the same drug, not only in their respective effective doses (*i.e*., *IC*_*50*_) but also in drug cooperativity (*i.e*., *n*) within 3D tumor models.

**Figure 6.**
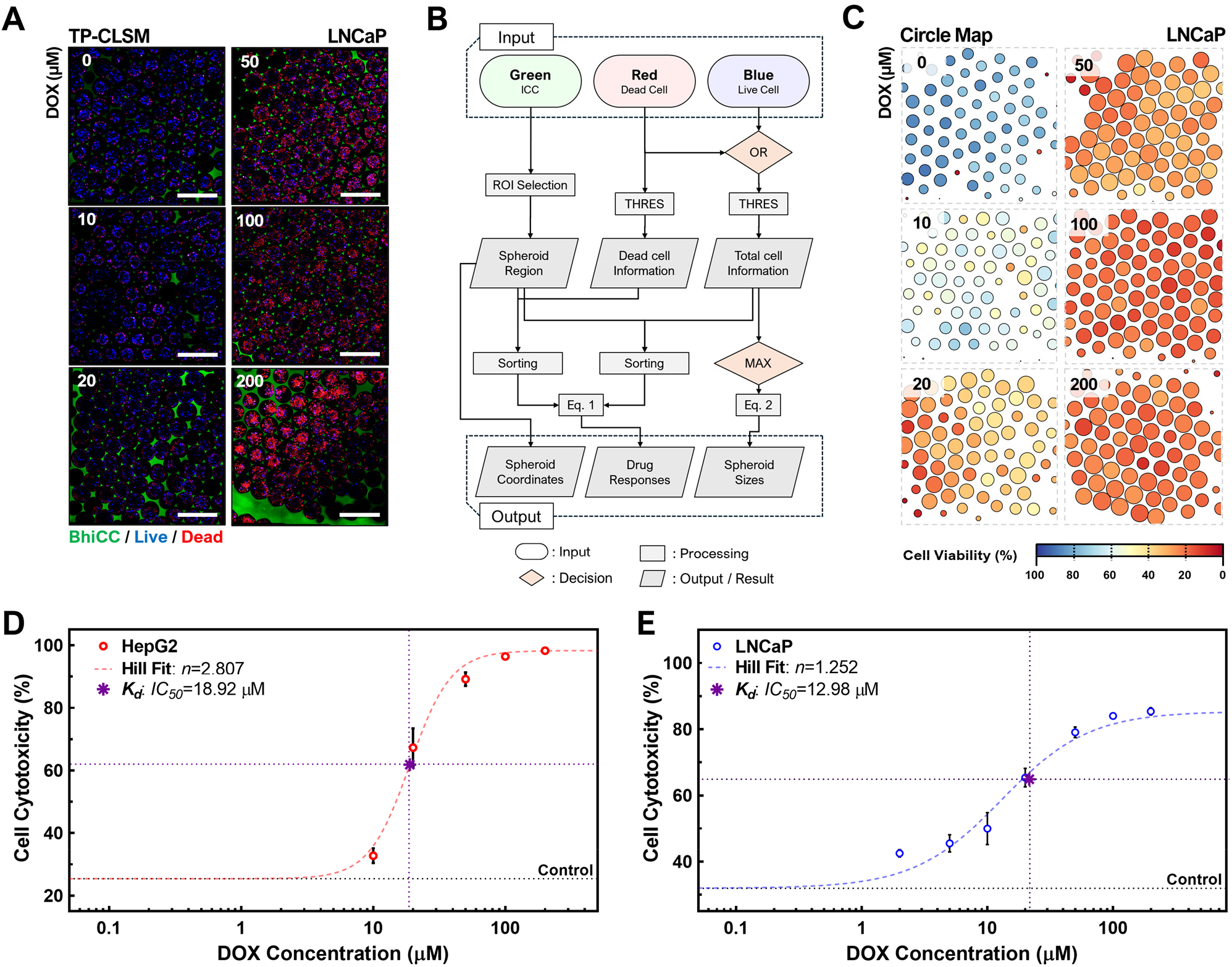
Proof-of-concept HC drug screening with BhiCC framework. **(A)** Single *Z*-plane TP-CLSM images of LNCaP spheroids within BhiCC framework after DOX treatment (24 h treatment and 24 h incubation; green: 5-DTAF on F-LMA, blue: live cells, and red: dead cells; scale bar: 500 µm). **(B)** Flow chart summarizing the automated image processing procedure developed. **(C)** Simplified circle mapping images of LNCaP spheroids within BhiCC framework after DOX treatment (24 h treatment and 24 h incubation). **(D-E)** Drug response curves generated by statistical collection from automated image processing from TP-CLSM images of **(D)** HepG2 and **(E)** LNCaP spheroids within BhiCC framework (error bar: ±SD; *N*=4 as number of images from each sample).

To advance implementation of quantitative analysis for HC screening on drug responses of cancer spheroids within the BhiCC framework, we designed an automated image processing method using TP-CLSM images (**Figure 6B**). Briefly, the void voxels from the green channel were employed to specify individual spheroid regions in the framework (*i.e*., region of interest; ROI), while blue and red voxels served as indicators of individual cell location and viability. Detailed elaboration on the automated image processing is provided in *Experimental Section*. Collecting and sorting the cell viability voxels enabled a more straightforward, comprehensive, and facile representation/analysis of the drug response from all sample groups. As an illustrative demonstration, we conducted circle mapping representing various information from the drug response TP-CLSM image: (1) Individual spheroid locations as circle coordinates, (2) individual spheroid sizes as circle diameters, and (3) individual spheroid drug response (spheroid viability) as circle colors (LNCaP: **Figure 6C** and HepG2: **Figure S17B**). Moreover, the image processing method we developed in this work was not only highly automated but also significantly simplify multi-channel 3D volumetric images rapidly (Response time: 0.4–0.8 sec per cellular spheroid), enabling the facile analysis and collection of entire data set from the drug response efficiently and effectively for HC drug screening (**Figure S18**). The statistical description of spheroid DOX response from the data processing is summarized in **Table S1**. To note, statistical values from all of the sample groups stemmed from 250–370 spheroids from 4 imaging replicates, indicating the HC potency of the BhiCC framework embedding high-yield biological replicates, increasing the statistical reliability of the system. Representatively, as shown in **Table S2**, we confirmed all of our samples showed statistical significance (*e.g*., most of the *P* values from comparison <0.0001 except two *P* values <0.001) while showing good-to-excellent accuracy (*e.g*., > 0.8) for the area under the curve (AUC) values in receiver operating characteristic (ROC) analysis.

This high statistical reliability enabled to plot of drug response curves from 3D spheroid structure with narrow SDs (LNCaP: **Figure 6E** and HepG2: **Figure 6D**; See **Table 1** for SDs). As a result, the qualitative observations from TP-CLSM images were clearly demonstrated quantitatively and well fitted with the Hill plots cell-respectively (*e.g*., the coefficient of determination was 0.9626 for HepG2 and 0.9368 for LNCaP). *IC*_*50*_ values from LNCaP were estimated at 12.98 μM while HepG2 showed 18.92 μM, elaborating on the differential sensitivity of the cellular spheroids to DOX and similar to the readouts from HT testing by the plate reader. These *IC*_*50*_ values are up to 15-to 50-fold higher compared to previous studies on DOX response in 2D cultured LNCaP and HepG2 cells, as well as the higher *IC*_*50*_ on 2D HepG2 than that of 2D LNCaP.^[76–82]^ The *IC*_*50*_s are higher than 2D culture, suggesting that the structural hinderance of drug transport, delivery, and effect within 3D tumorous tissue models is primarily due to the ECM expression. More importantly, in 3D imaging-based analysis, LNCaP showed a lower Hill coefficient *n* value than that of HepG2 (*e.g*., 1.252 and 2.807, respectively), indicating a more gradual dose-dependent response to drug treatment in LNCaP spheroids compared to HepG2, implying different drug adsorption and delivery to single cells in tumor tissue. This fact indicates the necessarily of HC drug response readout with 3D spatial information.

In summary, we presented a simple and straightforward methodology to create a BhiCC framework with precise dimensional control. Notably, we succeeded in fabricating the framework based only with nature-derived polymer-based hydrogels, alginate and agarose, which are advantageous on cost-effectiveness, safety, and ease in their utilization. Moreover, our approach on HCP assembling significantly streamlines the self-assembly process by employing the elasticity of alginate microgels forming precisely controllable interconnecting channels spontaneously, under mild conditions without using toxic chemicals, additional reactions, or complex equipment. This framework exhibits significant biological potency by simultaneously generating uniform, size-controlled, high-yield and physiologically reliable 3D *in-vitro* models (*e.g*., >30,000 spheroids per framework; ∼150 spheroids μL_-1_) which are scaffold-free and cancer-type independent. Furthermore, we validated its comprehensive application in both HT and HC drug screening as proof-of-concepts (*e.g*., adaptable with conventional HT drug screening; >20 spheroids mm^-2^ in HC drug screening), providing a promising platform for next-generation drug testing.

Automating fast HC screening, taking advantage of the uniform spheroids in HCP array offers comprehensive HC and HT drug testing capabilities, compared to conventional platforms that generate and analyze single spheroids per well. Beyond this proof-of-concepts, our system shows strong potential for incorporating more diverse readouts, such as gene expression and omics with spatial information, enabled by the high-yield, uniform spheroids and further development of the automated data processing algorithm for spatial multi-readouts will be promising to empower this HCHT platform. Moreover, numerous factors beyond ECM expression significantly influence drug efficacy in real tumors *in vivo*, including inherent heterogeneity in cell type and phenotype, diverse complexity in tumor microenvironments, pharmacokinetic-pharmacodynamic barriers in circulation, and variations in drug metabolism and clearance rates. Therefore, the systemic integration of the BhiCC framework with advanced systemic modeling techniques—such as pharmacokinetic conditions and pharmacodynamic barriers—is also essential. We will pursue follow-up studies to better reflect factors influencing drug efficacy and multi-readout analysis in real tumors, including co-culture systems or organoid cultures with stem cell-based approaches for greater physiological relevance, dynamic microfluidics to simulate pharmacokinetic conditions, patient-derived cell cultures to enhance patient-specific predictability, and ultimately, clinical validation of drug responses from the screened library.

## 3. Conclusion

In conclusion, we introduced a simple and efficient methodology for creating a BhiCC framework with precise dimensional control and applied them to 3D HC and HT drug screening. This innovative framework streamlined the fabrication process, enhancing biosafety and reducing procedural complexity while also improving the controllability of matrixed void spaces employing the innate elasticity of microgel template. The high crystallinity and uniformity in the resulting BhiCC framework led to highly parallelized support for generation of precisely size-controlled spheroids in the high-yield domain regardless of cancer cell type. The spheroids from the BhiCC framework demonstrated significant biological potency by producing ECM spontaneously, forming physiologically relevant scaffold-free cellular spheroids as 3D *in-vitro* models. Additionally, the spontaneous, uniform, high-yield spheroid generation within the HCP domain provided an ideal platform for developing and applying automated data processing, facilitating efficient cellular response collection, representation, and analysis. Our validation with proof-of-concept HT and HC drug screenings illustrate the framework’s comprehensive applicability, versatility, and potential for universal use in future works across various fields, including drug testing, screening, diagnostics, therapy development, and tissue engineering, expected to enhance the accuracy of predictive models for human responses and deepen the understanding of them.

## 4. Experimental Section

### Materials and Reagents

Alginic acid sodium salt (180947), calcium chloride dihydrate (CaCl_2_; 223506), sodium hydroxide anhydrous (NaOH; S5881), and alginate lyase powder (A1603) were purchased from Sigma-Aldrich (MO). Cryopres dimethyl sulfoxide (DMSO; 092780148) was purchased from MP Biomedicals (CA). Certified^™^ low melt agarose (LM agarose; 161-3111) was purchased from Bio-Rad (CA). 5-[(4,6-Dichlorotriazin-2-yl)amino]fluorescein hydrochloride (5-DTAF; 21811-74-5) was purchased from Chemodex (Switzerland). SylgardTM 184 silicone elastomer (polydimethylsiloxane, PDMS; 102092-312) was purchased from VWR (PA).

Minimum essential medium Eagle 1 X (MEM; 10-010-CV) and phosphate-buffered saline 1 X (PBS; 21-040-CM) were purchased from Corning (NY). Eagle’s Minimum Essential Medium with L-glutamine (EMEM; 30-2003) was purchased from American Type Culture Collection (ATCC; VA). RPMI-1640 Medium with L-glutamine (RPMI-1640; 112-023-101) was purchased from Quality Biological Inc (MD). Dulbecco’s Modified Eagle Medium/Nutrient Mixture F-12 (DMEM/F-12; 11320033) was purchased from Fisher Scientific (MA). Avantor Seradigm USDA-approved origin fetal bovine serum (FBS; 1300–500) was purchased from VWR (PA), kept at temperature of *™*20 °C. Antibiotic antimycotic (15240096) was purchased from Fisher Scientific (MA). Gibco™ Trypsin-EDTA (25200072) was purchased from Thermofisher Scientific (MA). 4% Paraformaldehyde in 0.1 M phosphate buffer (15735-50S) was purchased from Electron Microscopy Sciences (PA). CytoTrace™ Red CMTPX (22015) and Cell Explorer^™^ Live Cell Tracking Kit (22620) were purchased from AAT Bioquest (CA). LIVE/DEAD^™^ Cell Imaging Kit (488/570) (R37601) was purchased from Thermofisher Scientific (MA). Doxorubicin Hydrochloride (DOX; D4193) was purchased from Tokyo Chemical Industry Co. (Tokyo, Japan).

### Alginate Hydrogel Characterization and Microgel Fabrication

4 g of alginic acid were dissolved in 100 mL of deionized water (DIW), mixed for 1 h using a vortex mixer, and then autoclaved at 121 °C for 30 min. The resulting solution was diluted to final concentrations of 2.0, 2.5, and 3.0%_w/v_, followed by another 1 h of vortex mixing for uniform polymer solution formation. For the rheology test of alginate hydrogels, a hybrid rheometer (HR-2, TA Instruments; DE) was set with rotational geometry with 8 mm parallel plate. The gap was set at 800 μm and the hydrogel sample was located between the parallel plate and HR-2. A rheological strain sweep (*e.g*., 0.01-10% in rotational strain *ε*_*S*_ at a fixed *f* of 1 rad/s) was performed to collect compressive stress *σ*_*S*_ for each *ε*_*S*_. For the compression test of alginate hydrogels, the hybrid rheometer was set with compression fixture (disc) geometry with 8 mm parallel plate with 800 μm gap. A compressive oscillation sweep (*e.g*., 0.01-10% in compressive strain *ε*_*C*_ at a fixed *f* of 6.28 rad/s) was performed to collect compressive stress *σ*_*C*_ for each *ε*_*C*_. The compressive modulus *E* was determined from the linear fitting of the first 11 points in *σ*_*C*_-*ε*_*C*_ curves for each hydrogel.

For the electrospraying process, a commercially available apparatus, Spraybase_®_, was employed for (Spraybase; MA). A 26-gauge flat-end metal needle (S-0135-0001-01; Spraybase) was positioned at an injection distance of 210 mm from the spraying reservoir containing 30 mL of a 200 mM CaCl_2_ solution. Each alginic acid solution was then pumped through the metal needle at a pressure of 0.8 bar and sprayed into the spraying reservoir under a 20.0 kV DC voltage. The spraying operation was performed for 30 min as one cycle before alginate microgels were collected. For each cycle end, the entire system, including tubing and needle, underwent washing with deionized water three times. The resulting microgels were located on a 15 mm glass bottom dish and imaged with VWR Basic Inverted Microscope (76317-470, VWR; PA) with OMAX 5.0 MP High Sensitivity CCD Digital Camera (A3550UPA-R75, OMAX; WA), followed by a diameter estimation using ImageJ software.

### Agarose Hydrogel Fabrication and Characterization

To label the 5-DTAF dye into LM agarose, LM agarose and 5-DTAF underwent a reaction in a basic-conditioned solution serving as the reaction buffer. 5-DTAF powder was dissolved in DMSO to form a 10 mM solution and NaOH pellet was dissolved in DIW to form a 1 M solution. 250 mg of agarose powder was suspended in 8.55 mL of DIW for 10 min gentle mixing for hydration. The hydrated agarose solution was mixed with 500 µL of the 10 mM 5-DTAF solution for 5 min using a vortex mixer. Subsequently, 950 µL of the 1 M NaOH solution was slowly added dropwise. The resulting mixture was gently stirred at room temperature for 1.5 h, allowed to be stabilized for 10 min, and centrifuged (*e.g*., 1000 rpm for 3 min) to facilitate the separation of hydrated agarose and supernatant. The supernatant was carefully removed, leaving behind the hydrated agarose precipitate. The resulting precipitate was dispersed in 200 mL of DIW and precipitated for 10 min, before the supernatant removal. This washing step was repeated for three times. For fabricating LM agarose hydrogel, 1 g of agarose powder was suspended in 50 mL of DIW for 10 min for hydration. The mixture was then centrifuged (*e.g*., 1000 rpm for 3 min) to separate the hydrated agarose from the supernatant. The supernatant was removed, leaving the hydrated agarose precipitate. This un-labeled precipitate was mixed with the precipitate from the 5-DTAF labeling step in a 10%_w/w_ ratio (*e.g*., intact : labeled = 9 : 1 in weight ratio), cast into 50 mL conical tube. The resulting sample was heated at 70 °C for 30 min, followed by cooling in room temperature for 30 min. The resulting LM agarose hydrogel was sliced within 1.5 mm-thickness cylinder shape and re-shaped within the desired mold (*e.g*., a glass-bottom dish; 801002, NEST Scientific; NJ) by repeating the same temperature control above. The final agarose content inside the LM agarose hydrogel was estimated by comparing the weight of agarose powder *versus* the weight of the final hydrogel.

For the mechanical testing, 5-DTAF-tagged LM agarose hydrogel (*e.g*., 15.6%_w/v_) and alginate hydrogels (*e.g*., 2.0%, 2.5%, 3.0%_w/v_) were meticulously prepared in a cylindrical geometry, featuring a thickness of 0.8 mm and a diameter of 10 mm. For the viscoelasticity measurement, a dynamic oscillatory rheology test was performed with an HR-2 with 8 mm parallel plate geometry. The gap was set at 800 μm and the hydrogel sample was located between the parallel plate and HR-2. For each hydrogel, a frequency sweep (*e.g*., 0.1–10 rad/s in oscillation frequency *f* at a fixed shear strain *E*_*S*_ of 1%) was initially conducted to ensure operation within the linear viscoelastic regime. Subsequently, an amplitude sweep (*e.g*., 0.01–10% in *E*_*S*_ at a fixed *f* of 1 rad/s) was performed to collect storage modulus *G’* and loss modulus *G”*. The thermos-responsive feature of LMA hydrogel was confirmed by monitoring *G’* and *G”* under temperature sweep with a fixed *E*_*S*_ of 1% at a fixed *f* of 1 rad/s. For the fluorescence excitation-emission sweep test from 5-DTAF, the fluorescence emission property of resulting agarose hydrogel was measured both throughout a range of emission wavelengths (*e.g*., 515–600 nm; 5 nm intervals) with fixed excitation wavelength (*e.g*., 492 nm) and throughout a range of excitation wavelengths (*e.g*., 400–490 nm; 5 nm intervals) with a fixed emission wavelength (*e.g*., 516 nm) by a plate reader (Tecan Infinite 200Pro, Tecan; Switzerland). The uniform tagging of 5-DTAF throughout the hydrogel was verified with multi-photon imaging from a point scanning confocal microscope (Leica Stellaris 8 DIVE, Leica; Germany).

### BhiCC framework Fabrication

To form the microgel assembly, 6.0 × 10_4_ alginate microgel was prepared in 1 mL DIW. Subsequently, 250 µL of the microgel solution was deposited into the base of a glass-bottom dish and delicately covered with a cover glass. Using a bench-top sonicator (SU-743A, KECOOLKE; China), the glass-bottom dish underwent a 20-second sonication process, eliminating excess water from the glass-bottom part. This sonication process was repeated until no more water phase emerged. Finally, the cover glass was gently removed from the glass-bottom dish, leaving the HCP assembly of alginate microgels.

To cast the LM agarose hydrogel onto the HCP assembly, the glass-bottom dish-molded LM agarose hydrogel was positioned atop the HCP assembly in the glass-bottom dish, followed by putting a cover glass on top of the LM agarose hydrogel subsequently. The system was heated to 70 °C for 30 min, gently pressed to ensure filling LM agarose to all spaces in the glass-bottom dish, and then allowed to cool at room temperature for 30 min to solidify LM agarose hydrogel. Subsequently, the cover glass was gently removed, leaving behind the CC hydrogel containing the HCP assembly of alginate microgels encapsulated within the LM agarose hydrogel. To degrade the HCP assembly within the CC hydrogel, two sequential approaches were combined: enzymatic cleavage of alginic acid and chemical chelation of Ca^2+^ ions. To cleave the alginic acid chain, 1.5 mL of 1 mg*·*mL^-1^ alginate lyase was applied to the CC hydrogel at 37 °C for 3 h, followed by washing with DIW. The cleaved CC hydrogel was then exposed to 10 mL of PBS for 30 min, and the PBS was replaced 10 times until the hydrogel phase became transparent. The final BhiCC framework was washed with 2 mL PBS and underwent 30 min of UV sterilization. The formation of iCC structure inside the resulting hydrogel was verified with multi-photon imaging function of Leica Stellaris 8 DIVE Point Scanning Confocal Microscope (*i.e*., TP-CLSM), processed, and analyzed using ImageJ software.

### Cell Culture and Spheroid Generation in BhiCC Framework

Hepatocellular carcinoma human (HepG2; HB-8065) [American Type Culture Collection (ATCC), VA] cells were cultured in MEM supplemented with 10% FBS and 1% AA. Michigan Cancer Foundation-7 (MCF-7; 86012803) [Sigma-Aldrich, MO] cells were maintained in EMEM supplemented with 10% FBS and 1% AA. Lymph node carcinoma of the prostate (LNCaP; CRL-1740) [American Type Culture Collection (ATCC), VA] cells were cultured in RPMI 1640 supplemented with 10% FBS and 1% AA. Human neuroblastoma (SH-SY5Y; CRL-2266) [American Type Culture Collection (ATCC), VA] cells were cultured in DMEM/F-12 supplemented with 10% FBS and 1% AA. All cell lines were cultivated in a humidified incubator (MCO-15AC; Sanyo, Osaka, Japan) at 37 °C with 5% CO_2_ and routinely passaged to sustain exponential growth (*e.g*., Passages 6–12). When the cell lines reached > 80% confluency, they were washed with PBS, trypsinized using a trypsin-EDTA solution, and condensed into a 1 × 10^7^ mL^-1^ suspension for cell tracker staining or seeding into the BhiCC framework. To create the stock solution of the red cell tracker (*e.g*., CMTPX), 50 µg of CytoTrace^™^ Red CMTPX was dissolved in 7.28 µL of DMSO to form a 10 mM solution, which was then stored at -20 °C. For red cell tracker staining, 1 × 10^7^ mL^-1^ of trypsinized cells were treated with 5 µM of CMTPX within 2 mL of the cell culture medium for 1 h, followed by washing with PBS and resuspension into 1 × 10^7^ mL^-1^ in 1 mL fresh media. For blue cell tracker staining, 1 × 10^7^ mL^-1^ of trypsinized cells were treated with 20 µM of Cell Explorer™ Live Cell Tracking Kit within 2 mL of the cell culture medium for 30 min, followed by washing with PBS and resuspension into 1 × 10^7^ mL^-1^ in 1 mL fresh media.

To seed the HepG2, LNCaP, or SH-SY5Y cells into the BhiCC framework, 4.5 mL of a fresh cell culture medium and deposited on the BhiCC framework inside the perpendicularly-designed PDMS-molded 50 mL centrifuge tube. 1.5 × 10^7^ cells were suspended in 250 µL media and gently mixed with the 4.5 mL of the framework-embedding media in the centrifuge tube. The centrifuge tube was centrifuged at 400 rpm for 3 min followed by the gentle mixing of cell pellet outside the BhiCC framework. This centrifugation was repeated two times (*e.g*., Total 3.0 × 10^7^ cells per framework), followed by the final centrifugation with 400 rpm for 3 min. The BhiCC framework was moved into non-cell-treated 60 mm culture dish with 5 mL of media and incubated for 72 h at 37 °C with 5% CO_2_ followed by the media change for every 48 h. To seed the MCF-7 cells into the BhiCC framework, same procedure was applied with HepG2 and LNCaP case above, while 5.0 × 10^6^ cells was seeded on each centrifugation (*e.g*., Total 1.0 × 10^7^ cells per framework).

### Cellular Spheroid Characterization and Analysis

For TP-CLSM imaging, the BhiCC framework containing cellular spheroids were immersed in 2 mL PBS after being fixed with 2% PFA for 3 h and washed with PBS. Imaging was performed using the multi-photon imaging function of the Leica Stellaris 8 DIVE Point Scanning Confocal Microscope and analyzed with ImageJ software. To quantify the diameters of cellular spheroids within the BhiCC Framework, spatial information for each void space in the framework was gathered from the 5-DTAF channel, followed by collecting CMTPX signals from each void. The CMTPX channel was binarized using the Threshold function in ImageJ software to delineate the spheroid regions, which were then converted into diameters for each Z depth. The maximum diameter across all Z depths was recorded as the cellular spheroid diameter for each cell line (*e.g*., > 4 locations per cell line; > 100 total spheroids per cell line) and analyzed.

### Histological Sectioning, Staining, and Analysis of Cellular Spheroids

For histological sectioning, the frameworks containing cellular spheroids were immersed in 2 mL of PBS after being fixed with 2% PFA for 3 h and washed with PBS. BhiCC frameworks were processed in an automatic processor (TP1020, Leica; Germany) according to the procedures in the instruction manual. Following processing, the samples were paraffin-embedded and cut into 4–5 μm sections using a microtome (RM2125, Leica; Germany). These sections were then stained with hematoxylin and eosin (H&E) and Masson’s Trichrome (MTS). A modified MTS protocol was applied as follows: slides were deparaffinized and rehydrated through a series of alcohols ranging from 100% to 70%. After rinsing with distilled water, Biebrich scarlet-acid fuchsin was applied for 7 min. Slides were then rinsed and incubated in phosphomolybdic-phosphotungstic acid solution for 7 min, followed by immediate transfer to aniline blue for 10 min. Differentiation was performed briefly (approximately 1 min) using 1% acetic acid. Finally, the slides were rehydrated, cleared, and a coverslip was applied using resinous mounting media. The slides with H&E and MTS were imaged with bright field imaging function of All-in-One Fluorescence Microscope (BZ-X810, Keyence; Japan) and analyzed with ImageJ software.

### 3D HT and HC Drug Testing from Cellular Spheroids in BhiCC Framework

For drug response testing, blue cell tracker-stained HepG2 and LNCaP cells were prepared as spheroids within 5-DTAF-tagged BhiCC framework (HepG2: 6-day culture; LNCaP: 5-day culture). A range of DOX solutions was prepared (*e.g*., 0, 10, 20, 50, 100, and 200 μM) in fresh cell culture media for each cell lines with a final volume of 2 mL. The spheroids-embedding BhiCC frameworks were incubated with the DOX-embedding media for 24 h at 37 °C with 5% CO_2_. Following two-time PBS washing (2 mL each), the frameworks were further incubated with fresh cell culture media for 24 h at 37 °C with 5% CO_2_, followed by two more PBS washing (2 mL each). Subsequently, the BOBO-3 Iodide in Cell Explorer^™^ Live Cell Tracking Kit was dissolved in the fresh media for each cell lines and treated to spheroids-embedding BhiCC framework after drug treatment for 15 min. Finally, the frameworks were immersed in 2 mL of PBS after being fixed with 2% PFA for 3 h and washed with PBS.

For HT testing, the framework samples were cored with 5 mm Biopunch^®^ (15111-50, Ted Pella, Inc.; CA) and located in each well of Corning® 96 well NBS™ Microplate (Black flat-bottom; 3991, Corning; NY). The fluorescence measurements from the frameworks were performed with three channels by a plate reader (Tecan Infinite 200Pro): 5-DTAF with emission at 516 nm under 492 nm excitation, blue live cell tracker with emission at 450 nm under 350 nm excitation, and BOBO-3 Iodide with emission at 600 nm under 570 nm excitation. For the spatial plate reader reading, a well-scanning function from the plate reader (Tecan Infinite 200Pro) was applied (6-by-6 filled circle measurement per well). The dataset from each blue (*i.e*., live cell signal) and red (*i.e*., dead cell signal) were sorted only if they overlapped with green channel (*i.e*., BhiCC framework), and collected as *FI*_*blue*_ and *FI*_*red*_, pixel-respectively. The collected *FI*_*blue*_ and *FI*_*red*_ were reorganized and further processed using MATLAB software. To generate drug response curves, corrected *FI*_*red*_*/FI*_*blue*_ values were represented for each pixel converted from the raw intensity ratios from voxel as described in **Equation (5)**.

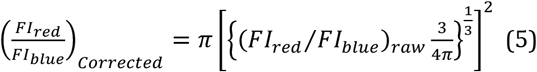

For HC testing, TP-CLSM imaging was performed using the multi-photon imaging function of the Leica Stellaris 8 DIVE Point Scanning Confocal Microscope. To facilitate quantitative analysis of drug-treated spheroids within the BhiCC framework, we developed an automated image processing method in Python, utilizing TP-CLSM images. Initially, the void spaces in the BhiCC framework were identified as regions of interest (ROIs) using the green channel of the images, pinpointing the coordinates of each void space. The red and blue channels were then merged (using an OR gate) and thresholded to delineate the 3D contours of the spheroids, independent of individual cell viability. Next, the red channel was thresholded and compared to the red-blue merged channel to assess cell viability within each 3D voxel. Finally, the viability data from each voxel were collected within the ROIs defined by the green channel, with each set of collected data representing the drug response of a corresponding spheroid in each ROI. These drug responses were then reorganized and further processed to generate drug response curves and circle maps. Detailed descriptions and instructions are available on GitHub (https://github.com/HyunsuJeon-ND/BhiCC-Framework-Drug-Testing).

## 5. Acknowledgements

We acknowledge funding from an American Cancer Society Institutional Research Grant (ACS IRG-17-182-04), the National Science Foundation Industry-University Cooperative Research Center (The Center for Bioanalytic Metrology), and NSF Career Award (NSF CBET-2337387). This research was funded in part by a Summer Graduate Research Fellowship and a Leiva Graduate Fellowship in Precision Medicine from the Berthiaume Institute for Precision Health at the University of Notre Dame. All the TP-CLSM imaging was carried out in part in the Notre Dame Integrated Imaging Facility, University of Notre Dame, using Leica Stellaris 8 DIVE Point Scanning Confocal Microscope. We thank Sara Cole and Sarah Chapman for the knowledge and expertise as well as time towards this research.

