## Supplementary material for "A Bioinert Hydrogel Framework for Precision 3D Cell Cultures: Advancing Automated High-Content and High-Throughput Drug Screening": SI

**This PDF file includes:**

- 2 Supplementary Notes
- 18 Supplementary Figures
- 2 Supplementary Tables

### Table of Contents

**Supplementary Note** ········ **S-2**

└ **Note 1.** Spherical Cap Formation in Microgel Assembly. ········ S-2

└ **Note 2.** Number of Spheroid Estimation in BhiCC Framework. ········ S-6

**Supplementary Figures** ········ **S-8**

└ **Figure S2.** Strain-sweep compressive analysis for alginate hydrogels. ········ S-8

└ **Figure S1.** Strain-sweep rheological analysis for alginate hydrogels. ········ S-8

└ **Figure S3.** Representation of alginate microgels from electrospraying. ········ S-9

└ **Figure S4.** Microgel assembly formation in a pseudo-isolated slit. ········ S-9

└ **Figure S5.** Fabrication process for LMA hydrogel. ········ S-10

└ **Figure S6.** Alginate-LMA merging process for CC hydrogel. ········ S-10

└ **Figure S7.** CC and BhiCC framework under microgel degradation. ········ S-11

└ **Figure S8.** TP-CLSM images of BhiCC framework. ········ S-11

└ **Figure S9.** Cell seeding and spheroid generation in BhiCC framework. ········ S-12

└ **Figure S10.** 3D reconstructed TP-CLSM images of cellular spheroid. ········ S-13

└ **Figure S11.** The spheroid area throughout Z positions in BhiCC framework. ········ S-14

└ **Figure S12.** Histological results for spheroid inside the BhiCC framework. ········ S-15

└ **Figure S13.** Two-color scatter mapping results from MTS. ········ S-16

└ **Figure S14.** Drug response results from single-point readout. ········ S-17

└ **Figure S15.** Heatmap of live cell fluorescence intensity from BhiCC framework. ········ S-18

└ **Figure S16.** Histogram results of BhiCC framework-embedded spheroids········ S-19

└ **Figure S17.** Proof-of-concept HC drug screening for HepG2. ········ S-20

└ **Figure S18.** Full image set of circle mapping images. ········ S-21

**Supplementary Tables** ········ **S-22**

└ **Table S1.** Statistical description for HC drug screening with BhiCC framework. ········ S-22

└ **Table S2.** The P-values and AUC values collected from datapoint comparison. ········ S-23

### Supplementary Notes

#### Note 1. Spherical Cap Formation in Microgel Assembly

(i) Single Microgel Standing on Solid Bottom

This description pertains to the estimation of channel formation in alginate microgel-based colloidal crystal (CC) assemblies. In this system, our microgels are in an aqueous phase and tend to sink to the bottom due to their density (*e.g.*, $\rho_{alg} = \rho_w(1 + C_{alg}) > \rho_w$ where $\rho_{alg}$: density of microgel, $C_{alg}$: alginic acid contents, and $\rho_w$: density of water). If the microgels are sufficiently soft to form channels with one another upon piling up, without disrupting their structure, the compression exerted by neighboring microgels should counterbalance the gravitational sinking, thus stabilizing the system in a steady state. We can start modeling this system mathematically, from the situation of a single microgel standing on solid bottom as the simplest case.

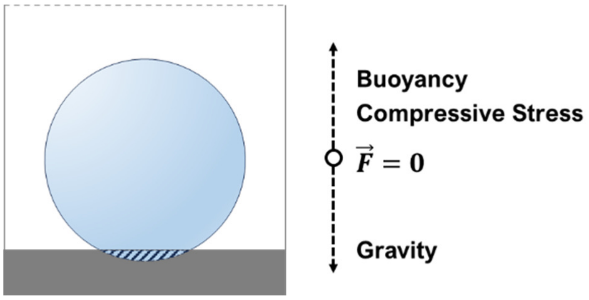


Since we assume that the microgels are sufficiently soft to form channels, they will take on a compressed shape resembling a spherical segment (*e.g.*, the non-dashed area in the drawing) without the spherical cap (*e.g.*, the dashed area in the drawing). This shape results from the balance between effective gravity and compression forces. We can consider the force balance involving compressive stress (*i.e.*, $\boldsymbol{F}_c$), buoyancy (*i.e.*, $\boldsymbol{F}_b$), and gravitational force (*i.e.*, $\boldsymbol{F}_g$) as **Equation (1)**.

$$\boldsymbol{F}_{net} = \boldsymbol{F}_c + \boldsymbol{F}_b + \boldsymbol{F}_g = 0 \quad \cdots \quad (\mathbf{1})$$

Here, we can describe $\boldsymbol{F}_g$ and $\boldsymbol{F}_b$ from simple physics where $r$ is radius of microgel and $g$ is gravitational acceleration as follows:

$$\boldsymbol{F}_g = \frac{4}{3}\pi r^3 \rho_w (1 + C_{alg}) g = -\boldsymbol{F}_b + \frac{4}{3}\pi r^3 \rho_w C_{alg} g$$

If the compression of microgel is not so severe thus falls to the linear elastic region in compressive strain-stress curve (See **Figure S2**), we can correlate compressive stress (*i.e.*, $\sigma_c$) and strain (*i.e.*, $\varepsilon_c$) with compressive modulus (*i.e.*, $E_c$) as follows: $E_c = \sigma_c / \varepsilon_c$. Accordingly, we can describe how much stress is generated under the compression of single microgel where $r'$ is the distance between the microgel center and the spherical cap plane, $S$ is the surface of the plane on the spherical cap as follows:

$$F_c = -\sum_S \sigma_c = -\oiint_S \sigma_c \, dA = -\oiint_S E_c \varepsilon_c \, dA = -\oiint_S E_c \frac{(r-r')}{r} \, dA = -E_c \frac{(r-r')}{r} \int_{r'}^{r} \pi (r^2 - x^2) \, dx$$
$$= -\pi \frac{(r-r')}{r} \left( \frac{2}{3} r^3 - r^2 r' + \frac{1}{3} r'^3 \right) E_c$$

Combining three component of force balance $\boldsymbol{F}_{net} = \boldsymbol{F}_c + \boldsymbol{F}_b + \boldsymbol{F}_g = 0$ gives us an equation describing correlation between the deformed radius of microgel (*e.g.*, $r'$) and $E_c$ as **Equation (3)** where the distance between the microgel center $Di = 2r'$ and the diameter of microgel $R = 2r$:

$$\boldsymbol{F}_{net} = -\pi \frac{(r-r')}{r} \left( \frac{2}{3} r^3 - r^2 r' + \frac{1}{3} r'^3 \right) E_c - \frac{4}{3} \pi r^3 \rho_w g + \frac{4}{3} \pi r^3 \rho_w (1 + C_{alg}) g = 0$$
$$r'^4 + (-r)\, r'^3 + (-3\, r^2)\, r'^2 + (5\, r^3)\, r' - 2r^4 \left( 1 - \frac{2\, \rho_w\, C_{alg}\, g}{E_c} \right) = 0$$

$$\left(\frac{Di}{2}\right)^4 - \left(\frac{R}{2}\right)\left(\frac{Di}{2}\right)^3 - 3\left(\frac{R}{2}\right)^2\left(\frac{Di}{2}\right)^2 + 5\left(\frac{R}{2}\right)^3\left(\frac{Di}{2}\right) - 2\left(\frac{R}{2}\right)^4\left(1 - \frac{2\, \rho_w C_{alg} g}{E_{c,alg}}\right) = 0 \quad \cdots \quad (\mathbf{2})$$

<u>(ii) Multiple Microgels Piling in Hexagonal Closed Packing (HCP)</u>

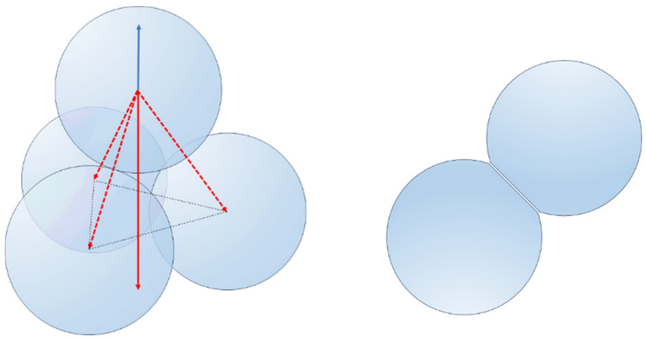

Similarly, we can describe the situation of multiple microgels standing by themselves. Since microgels will form HCP structures by themselves after dehydration, we can describe the force balance with the force balance as we did in single microgel standing. Notably, as shown in drawing below, $\boldsymbol{F}_g$ and $\boldsymbol{F}_b$ terms would not change in their absolute values while different values. Specifically, their summation will be changed since three microgels will support one microgel above. Hence, we project 1/3 on $\boldsymbol{F}_g + \boldsymbol{F}_b$ term in **Equation (1)**. On the other hand, $\boldsymbol{F}_c$ term need to be doubled, since now two deformed parts are compensating each other in this multiple microgel packing case. Notably, the weight of the microgel above third layer would be compensated with the microgel deformation on their side. Therefore, our force balance will be same as following **Equation (3)** where the distance between the microgel center $Di = 2r'$ and the diameter of microgel $R = 2r$:

$$\boldsymbol{F}_{net} = 2\boldsymbol{F}_c + \frac{1}{3}(\boldsymbol{F}_b + \boldsymbol{F}_g) = 0$$

$$\boldsymbol{F}_{net} = 2\pi\frac{(r-r')}{r}\left(\frac{2}{3}r^3 - r^2r' + \frac{1}{3}r'^3\right)E_c + \frac{1}{3}\left(\frac{4}{3}\pi r^3\rho_w g - \frac{4}{3}\pi r^3\rho_w(1+C_{alg})g\right) = 0$$

$$(+1)\,r'^4 + (-r)\,r'^3 + (-3r^2)r'^2 + (+5r^3)r' + \left(-2r^4\left(1 - \frac{\rho_w C_{alg} g}{3E_c}\right)\right) = 0$$

$$\left(\frac{Di}{2}\right)^4 - \left(\frac{R}{2}\right)\left(\frac{Di}{2}\right)^3 - 3\left(\frac{R}{2}\right)^2\left(\frac{Di}{2}\right)^2 + 5\left(\frac{R}{2}\right)^3\left(\frac{Di}{2}\right) - 2\left(\frac{R}{2}\right)^4\left(1 - \frac{\rho_w C_{alg} g}{3E_{c,alg}}\right) = 0 \quad \cdots \quad (\mathbf{3})$$

(iii) Channel Diameter Estimation

From the mathematical description in **S1.1** (i)-(ii), we can estimate the channel diameter from the microgel overlapping, eventually forming interconnecting channels in the resulting bioinert hydrogel-based iCC framework (BhiCC framework) with a following relationship where $d_c$ is channel diameter:

$$d_c = 2\sqrt{r^2 - r'^2} = 2\sqrt{\left(\frac{R}{2}\right)^2 - \left(\frac{Di}{2}\right)^2} \quad \cdots \quad (\mathbf{4})$$

Combining all, we provided a simple MATLAB script (*e.g.*, **MATLAB Script #1** below) which solve Equation (2) and Equation (3) to get real, positive, smaller-than-microgel-diameter channel diameters (*i.e.*, $C_d$) from respective $E_{c,alg}$ values. Notably, the solution from single microgel standing (*e.g.*, **Equation (2)**) can only be applied to the first layer of HCP assembly of microgel from bottom (*i.e.*, $n_{HCP} = 1$) while the multiple microgel solution (*e.g.*, **Equation (3)**) can be applied to upper layers ((*i.e.*, $n_{HCP} > 1$). The resulting plots are represented below. Notably, measured $E_c$ values for the microgels in this work ranged in 500-2000 kPa, which is expected to form 10-25 μm interconnecting channels theoretically.

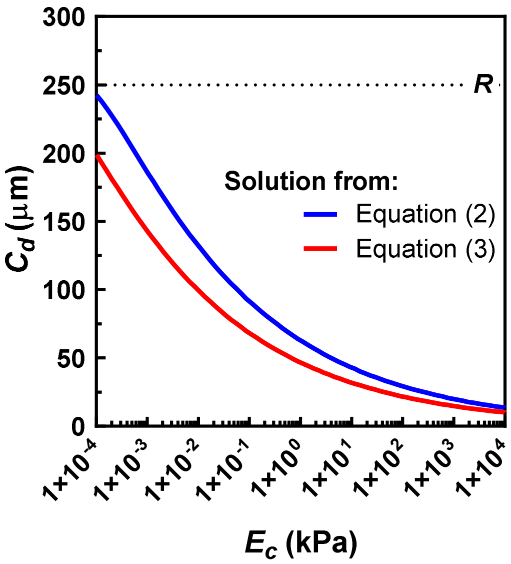

**MATLAB Code 1**: MATLAB Script

```
% Parameter input
R = 0.25;                % Microgel Diameter [mm]
Rho_w = 1e-4;            % Water density [g/mm^3]
g = 9806.65;             % Gravitational acceleration [mm/s^2]
C_alg = 0.025;           % Alginate content (2.5%)
E_c = 10.^linspace(-2, 8, 100);  % Compressive Modulus [g/mm/s^2]

% Preallocate result arrays
Di_SM_collect = zeros(1, length(E_c));
Di_MM_collect = zeros(1, length(E_c));

% Coefficients common to both equations
coeff_common = [(1/2)^4, -(R/2)*(1/2)^3, -3*(R/2)^2*(1/2)^2, 5*(R/2)^3*(1/2)];

% Solve Equation (2) and (3) in a single loop
for i = 1:length(E_c)
    % Equation (2)
    p_SM = [coeff_common, -2*(R/2)^4 * (1 - 2*Rho_w*C_alg*g/E_c(i))];
    r_SM = roots(p_SM);
    Di_SM = max(r_SM(real(r_SM) > 0 & imag(r_SM) == 0 & real(r_SM) < R), [], 'omitnan');
    if isempty(Di_SM), Di_SM = 0; end
    Di_SM_collect(i) = Di_SM;

    % Equation (3)
    p_MM = [coeff_common, -2*(R/2)^4 * (1 - Rho_w*C_alg*g/E_c(i)/3)];
    r_MM = roots(p_MM);
    Di_MM = max(r_MM(real(r_MM) > 0 & imag(r_MM) == 0 & real(r_MM) < R), [], 'omitnan');
    if isempty(Di_MM), Di_MM = 0; end
    Di_MM_collect(i) = Di_MM;
end

% Result Output
E_c = E_c';
Cd_SM = 2 * sqrt((R/2)^2 - (Di_SM_collect/2).^2)'; % Single Microgel Standing on Solid
Bottom
Cd_MM = 2 * sqrt((R/2)^2 - (Di_MM_collect/2).^2)'; % Multiple Microgels Piling

% Display Results
disp('Compressive Modulus (E_c):');disp(E_c); %[g/mm/s^2]
disp('Cd_SM (Single Microgel):');disp(Cd_SM); %[mm]
disp('Cd_MM (Multiple Microgels):');disp(Cd_MM); %[mm]
```

#### Note 2. Number of Spheroid Estimation in BhiCC Framework

This description explains how many microgels can be packed inside a given mold (*e.g.*, a 15 mm glass-bottom dish). To estimate the microgel packing, we make two simple assumptions: (1) the microgels form hexagonal close packing (HCP) during the time of interest, and (2) the microgels possess sufficient mechanical stiffness to achieve perfect elastic packing, with no deformation. With these assumptions, the problem can be simplified into a 2D circle packing problem within a larger circle, following HCP crystallinity. To determine the number of smaller circles in the HCP domain within a larger circle, we provide a simple MATLAB function (See **MATLAB Script #2** below). This function generates an HCP array of smaller circles from the center of the larger circle, checks whether their periphery falls inside the larger circle, and collects the coordinates of those that do. Notably, the HCP domain of small circles within a larger circle emerges from two different starting points: a single circle (single point (SP) layer) and three circles (triple point (TP) layer). When filling the microgel inside the cylindrical mold, the SP and TP layers alternate to form the HCP domain in a 3D cylindrical coordinate system. For instance, a 5-layer microgel assembly with HCP crystallinity will have two possible total microgel numbers ($N_{total}$): $N_{total} = 3N_{SP} + 2N_{TP}$ or $N_{total} = 2N_{SP} + 3N_{TP}$, depending on the starting point of the bottommost layer. The following plot shows the dependency of the packed microgel number on the mold diameter, assuming a 5-layer HCP domain.

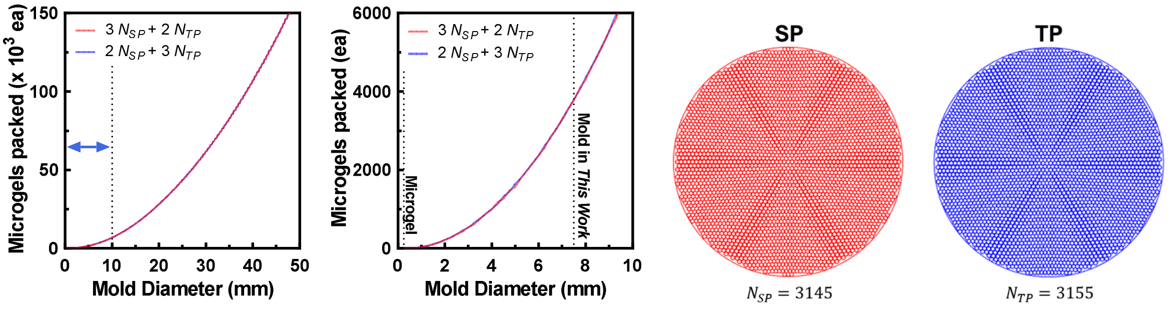


The illustration above represents a more specific example using a 15 mm glass-bottom dish as the mold for microgel packing. The estimated $N_{SP}$ was 3145, and $N_{TP}$ was 3155, resulting in a total microgel number ($N_{total}$) of about 15745-15755. Correlating this result with the high-yield spheroid/organoid generation feature of our framework, $N_{total}$ corresponds to the theoretical number of spheroids/organoids formed within a single framework.

**MATLAB Code 2**: MATLAB Function

```
function [FinalSP,FinalTP,XYSP,XYTP] = FindHCP(Iradius,Oradius)
%Number of circle in Hexagon packing fitted in Large circle
%Iradius represents radius of smaller circle
%Oradius represents radius of larger circle

%%HCP Domain Generation
base=Iradius*2;
X=[0]; Y=[0]; XYSP=[]; XYTP=[];
    for num=1:fix(Oradius/Iradius)/2+5
        x=zeros(num*6,1); y=zeros(num*6,1);
        x(1:6)=base*num*cos(2*pi/6.*(0:5));
```

```
        y(1:6)=base*num*sin(2*pi/6.*(0:5));
    if num>1
        for q=1:num-1
            start_x=x(2)-q*base;
            radi0=sqrt(start_x^2+y(2)^2);
            start_alpha=1/3*pi+pi*1/3*1/(num)*q;
            x(q*6+1:q*6+6)=radi0*cos(start_alpha+pi/3.*(1:6));
            y(q*6+1:q*6+6)=radi0*sin(start_alpha+pi/3.*(1:6));
        end
    end
        X=[X; x]; Y=[Y; y];
    end

%% If starts with single circle
FinalSP=0; figure(1)
set(gcf,'position',[10,10,500,500]);hold on;
xlim([-1.2*Oradius 1.2*Oradius]); ylim([-1.2*Oradius 1.2*Oradius]);
for i=1:length(X)
    Distance=norm([0;0]-[Y(i);X(i)]);
    if Distance<(Oradius-Iradius)
        XYSP=[XYSP;[X(i),Y(i)]];
        viscircles([X(i),Y(i)],Iradius,'Color','r','LineWidth',0.1);hold on;
        FinalSP=FinalSP+1;
    end
end
circle(0,0,Oradius,'r-');axis equal;

%% If starts with three circles (triangle shape)
FinalTP=0;figure(2)
set(gcf,'position',[10,10,500,500]);hold on;
xlim([-1.2*Oradius 1.2*Oradius]);ylim([-1.2*Oradius 1.2*Oradius]);
for i=1:length(X)
    Distance=norm([Iradius/2;Iradius]-[Y(i);X(i)]);
    if Distance<(Oradius-Iradius)
        XYTP=[XYTP;[X(i),Y(i)]];
        viscircles([X(i),Y(i)],Iradius,'Color','b','LineWidth',0.1);hold on;
        FinalTP=FinalTP+1;
    end
end
circle(Iradius,Iradius/2,Oradius,'b-');axis equal;
end
```

### Supplementary Figures

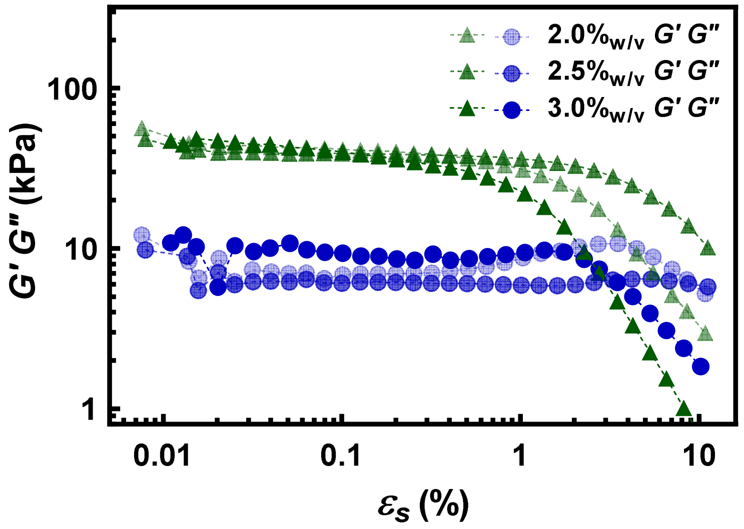


**Figure S1**. Result of strain-sweep rheological analysis for alginate hydrogels from different alginic acid contents ($C_{alg}$; 2.0-3.0%$_{w/v}$). The measurements were performed with 0.01-10% in rotational strain $\varepsilon_s$ at a fixed $f$ of 1 rad/s.

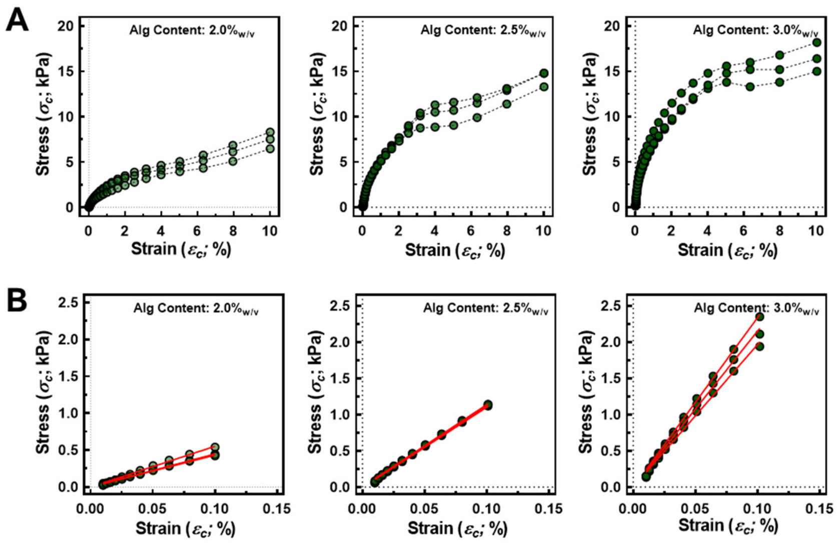


**Figure S2**. Result of strain-sweep compressive analysis for alginate hydrogels from different $C_{alg}$ (2.0-3.0%$_{w/v}$). The measurements were performed with 0.01-10% in compressive strain $\varepsilon_c$ at a fixed $f$ of 6.28 rad/s. **(A)** Plots showing full range of $\varepsilon_c$. **(B)** Plots showing linear $\sigma_c$-$\varepsilon_c$ range and linear regressions.

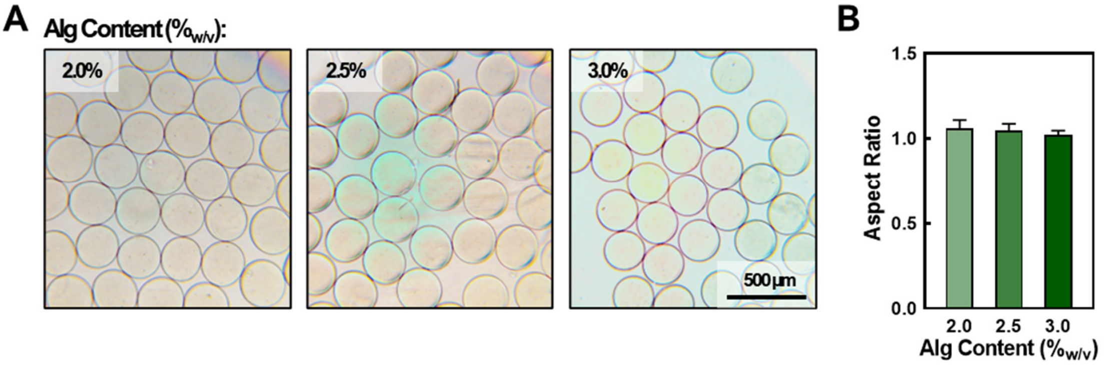


**Figure S3**. Representation of alginate microgels from different $C_{alg}$ (2.0-3.0%$_{w/v}$) fabricated by electrospraying. **(A)** Representative microscopic images of microgels from each $C_{alg}$ (left: 2.0%$_{w/v}$, middle: 2.5%$_{w/v}$, right: 3.0%$_{w/v}$; scale bar: 500 µm). **(B)** Aspect rationes for all microgel groups. (error bar: ±SD; *N*=60 or more).

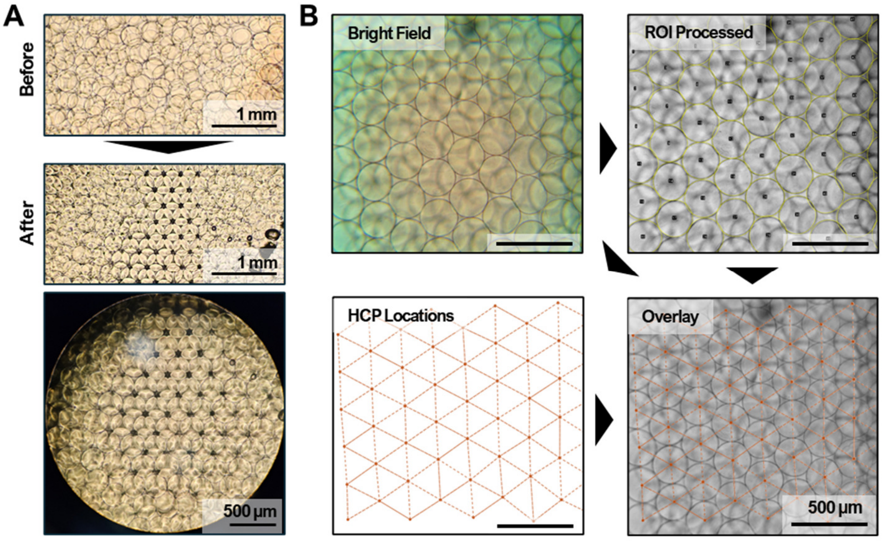


**Figure S4**. Microgel assembly formation in a pseudo-isolated slit. **(A)** Digital images of microgels before (top) and after (middle and bottom) the sonication process (scale bar: 1 mm). **(B)** Representative microscopic image of the microgel assembly and the demonstration procedure for its hexagonal close-packed (HCP) structure (scale bar: 500 µm).

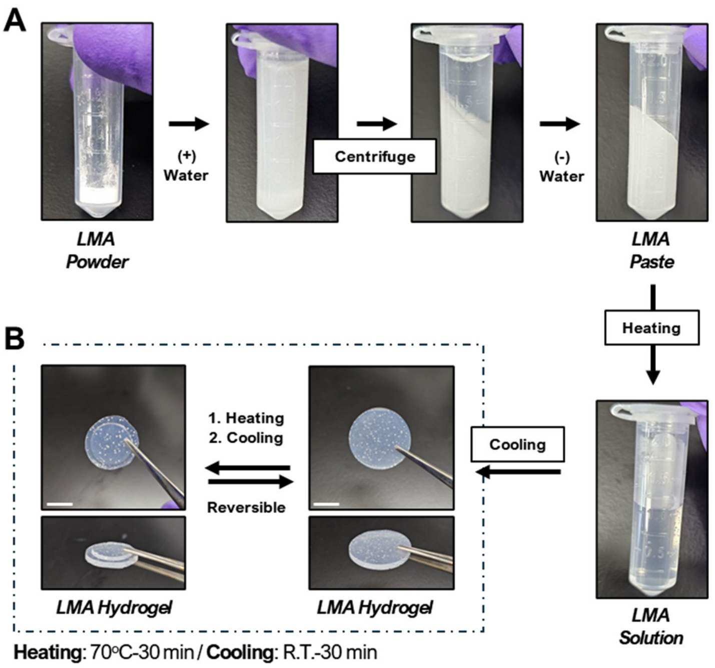


**Figure S5.** Fabrication process and moldability demonstration for low-melting agarose (LMA) hydrogel. **(A)** Digital images of the product from the fabrication process of LMA hydrogel, including hydration, isolation, heating, and cooling. **(B)** Digital images showing the thermo-responsive moldability of LMA hydrogel.

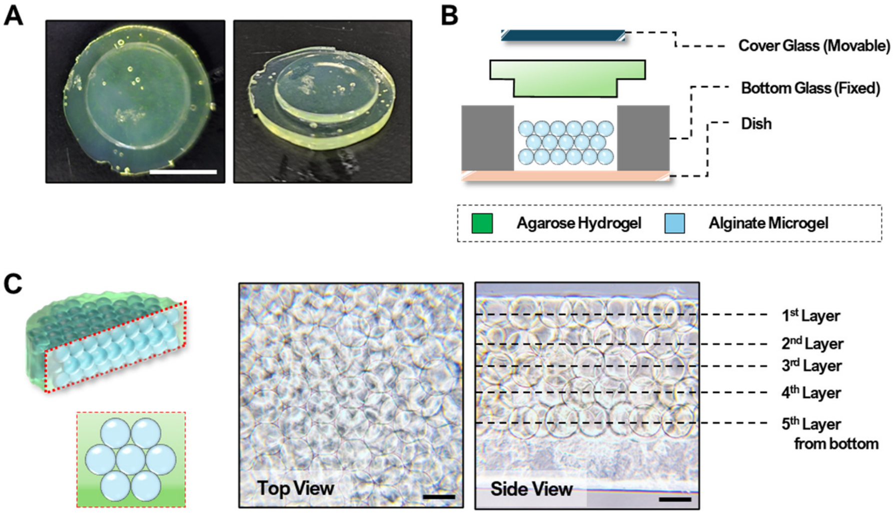


**Figure S6.** Representative images and schemes illustrating alginate-LMA merging process, resulting in a colloidal crystal (CC) hydrogel. **(A)** Digital images of a glass-bottom dish-molded fluorescent LMA (F-LMA) hydrogel (scale bar: 1 cm). **(B)** Schematic illustration of alginate microgel-LMA hydrogel merging process. **(C)** Illustration (left) and representative microscopic images of resulting CC hydrogel (center: top-view microscopic image and right: side-view microscopic image after sectioning; scale bar: 200 μm).

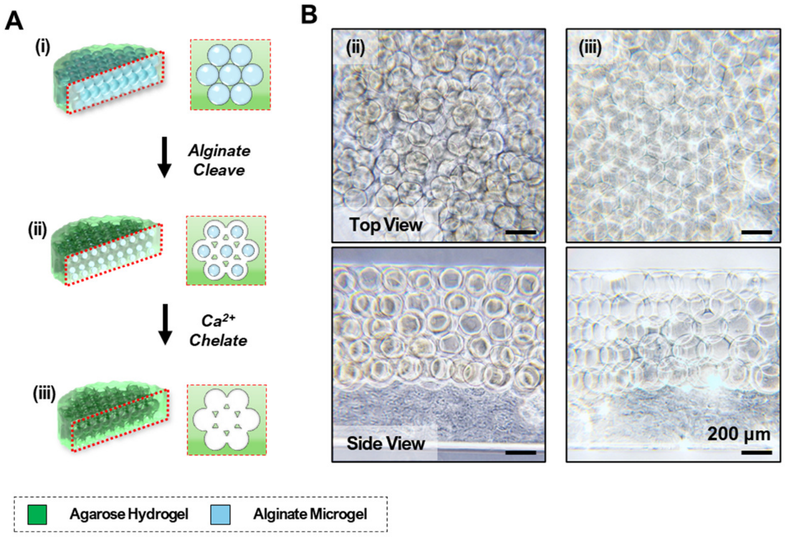


**Figure S7**. Schematic illustration and microscopic images of CC and BhiCC framework under alginate microgel degradation process. **(A)** Schematic illustration for **(i)** the CC hydrogel, **(ii)** the cleaved CC hydrogel after alginate lyase treatment, and **(iii)** the resulting ICC hydrogel after $Ca^{2+}$ chelation. **(B)** Microscopic images of **(ii)** the cleaved CC hydrogel (left) and **(iii)** the resulting BhiCC framework (right; scale bar: 200 μm).

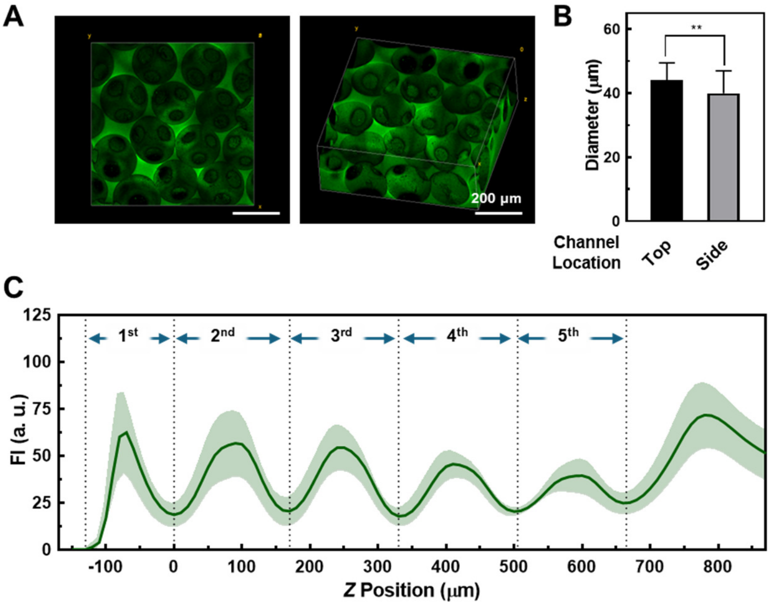


**Figure S8.** Representative two-photon confocal laser scanning microscopic (TP-CLSM) image of resulting BhiCC framework and its analysis. **(A)** 3D reconstructed image of BhiCC framework from TP-CLSM (green: 5-DTAF on LMA hydrogel; Scale bar: 200 μm). **(B)** Channel diameters on the hydrogel (black bar; top channels) and in the hydrogel (gray bar: side channels) (error bar:

±SD; $N = 5$; **: $P < 0.01$). **(C)** Plot of averaged fluorescent intensities (FI) *versus* $Z$ positions of each $Z$ stack image in BhiCC framework (green: 5-DTAF on LMA hydrogel).

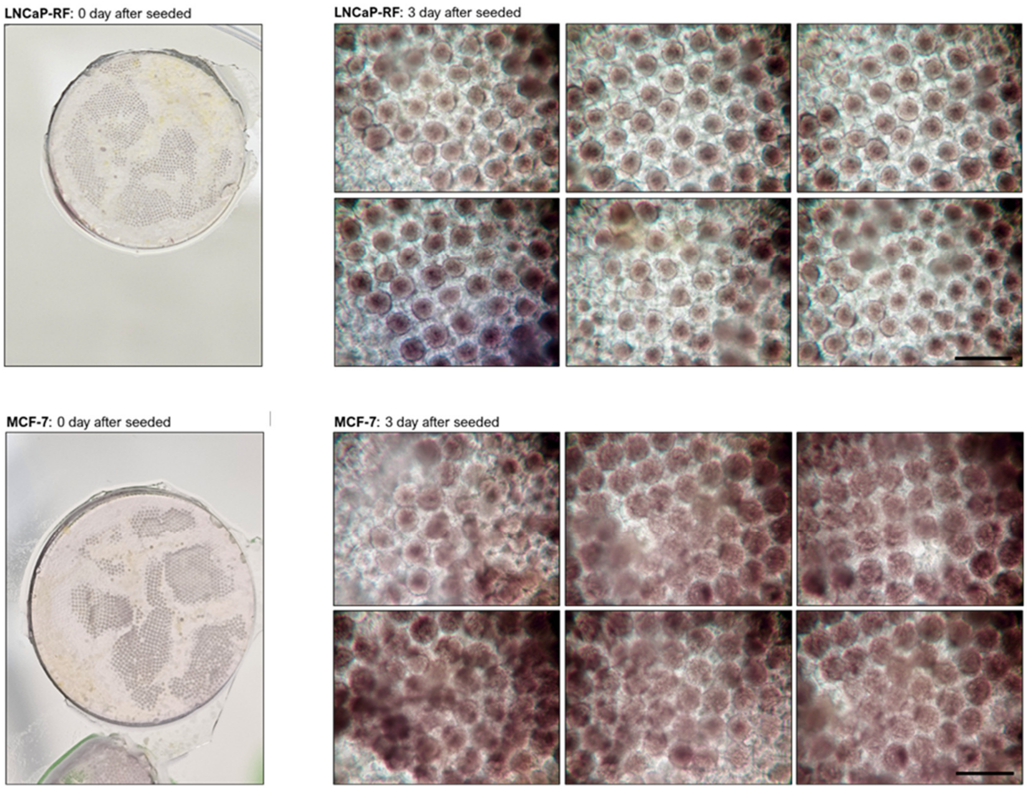


**Figure S9.** Digital image showing cell seeded inside the BhiCC framework (*i.e.*, after 0 h) and microscopic images showing spheroid generation inside the BhiCC framework (*e.g.*, after 72 h; Scale bar: 500 μm). The images shown as representatives for human prostate carcinoma (*i.e.*, LNCaP, top) and human adenocarcinoma (*i.e.*, MCF-7, bottom).

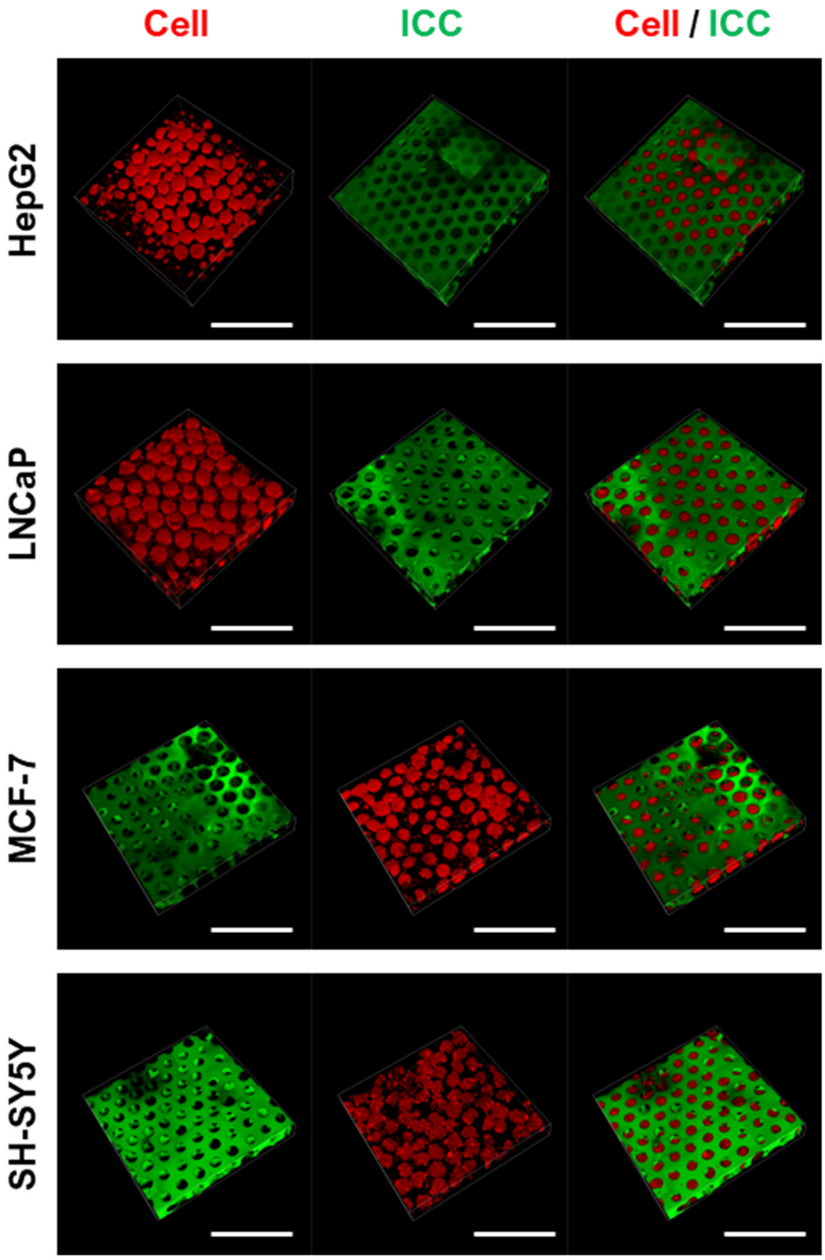


**Figure S10.** 3D reconstructed TP-CLSM images showing cellular spheroid formation inside the BhiCC framework regardless of cell lines (green: 5-DTAF on LMA; red: CMTPX live cell tracker; first row: HepG2, second row: LNCaP, third row: MCF-7, and fourth row: SH-SY5Y; scale bar: 500 μm).

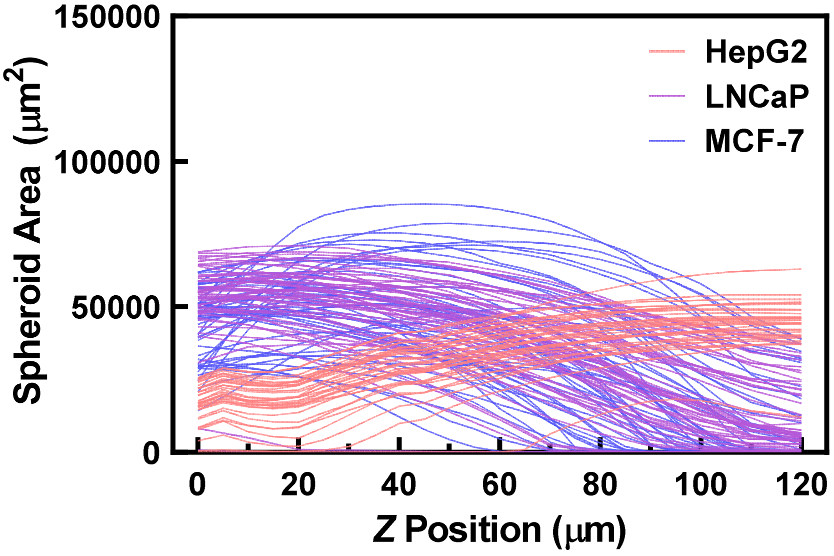


**Figure S11.** The spheroid area throughout *Z* positions of each *Z* stack image in BhiCC framework collected from TP-CLSM images. The plot color represents HepG2 in red, LNCaP in purple, and MCF-7 in blue color.

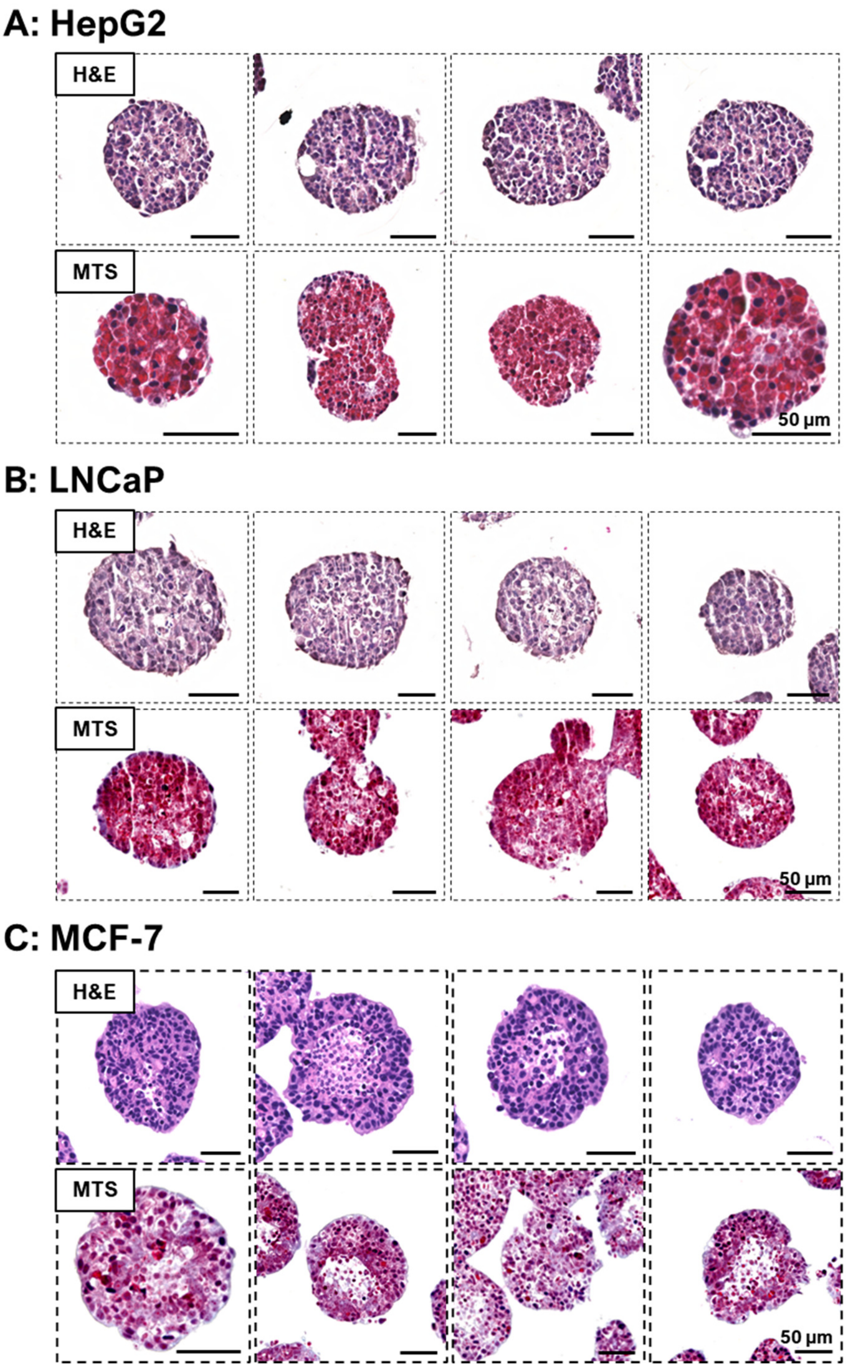


**Figure S12.** Histological results for spheroid inside the BhiCC framework stained with Hematoxylin and eosin staining (*i.e.*, H&E; Top of each panel) and Masson's Trichrome staining (*i.e.*, MTS; Bottom of each panel). **(A)** HepG2, **(B)** LNCaP, and **(C)** MCF-7 spheroids were represented (Scale bar: 50 μm).

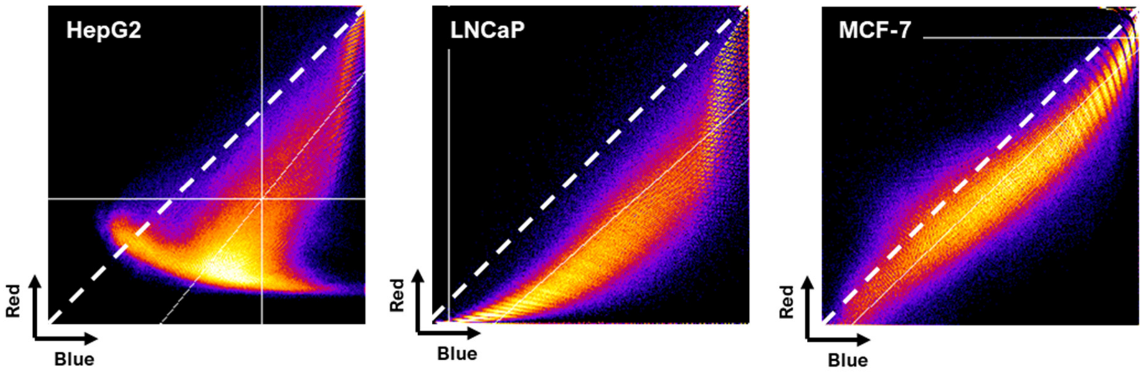


**Figure S13.** Two-color scatter mapping results from HepG2 (left), LNCaP (center), and MCF-7 (right) spheroid from blue and red color represented in MTS.

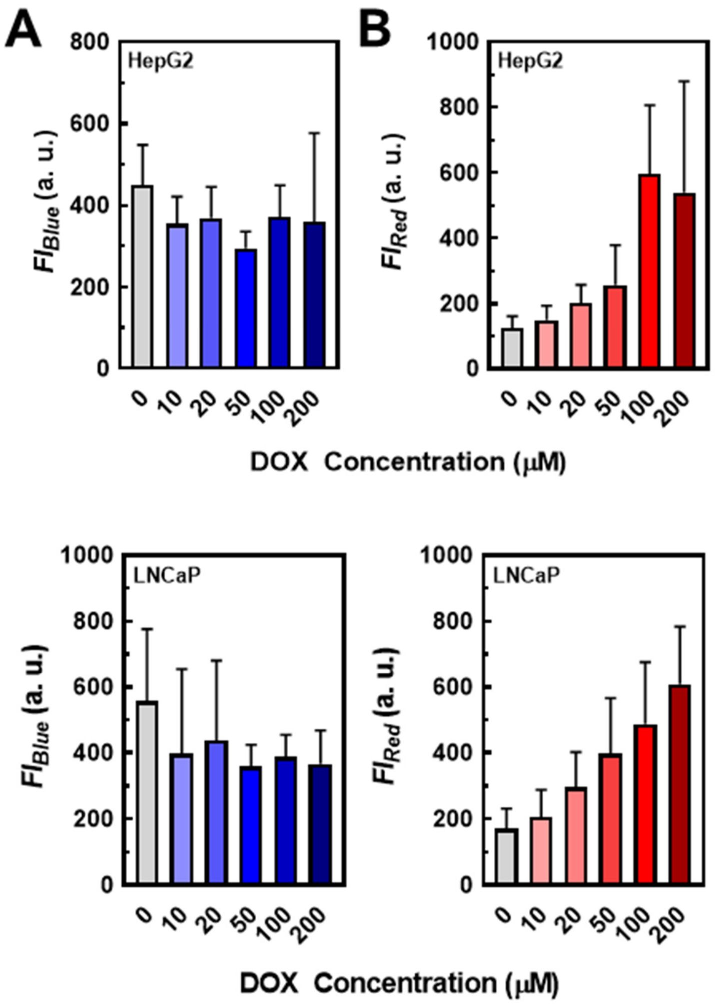


**Figure S14.** Drug response results of BhiCC framework-embedded spheroids from (**A)** HepG2 and **(B)** LNCaP as single point readout by the plate reader. Left plots of each panel represents the live cell signal (*i.e.*, $FI_{blue}$) from spheroids while right plots indicating dead cell signal (*i.e.*, $FI_{red}$) from drug response (error bar: ±SD; $N$ = 4).

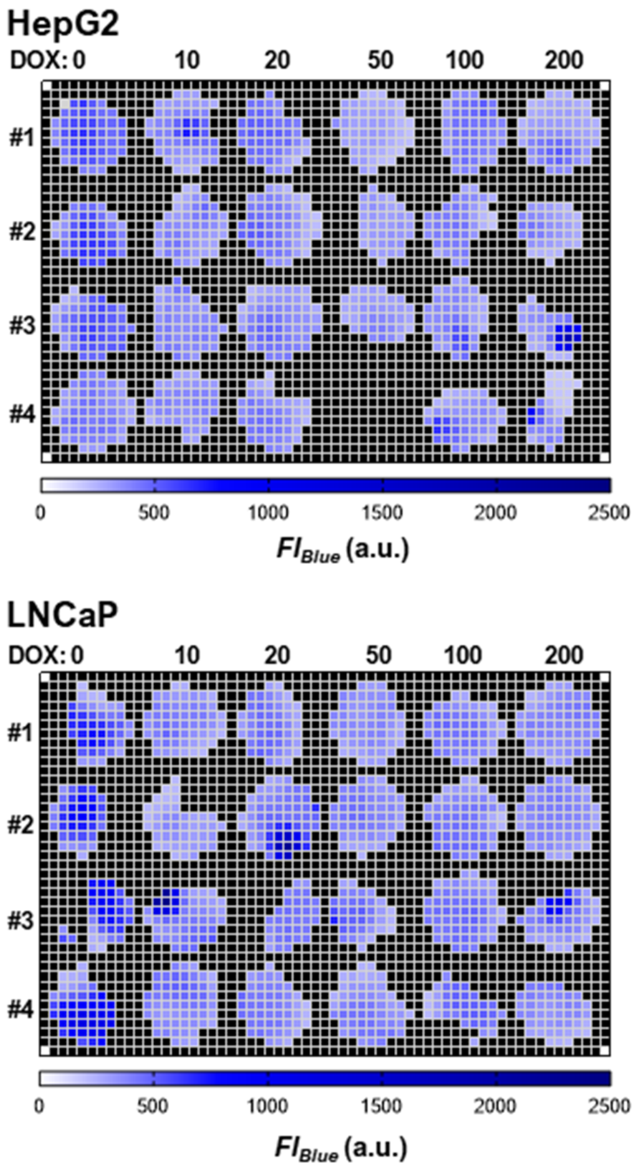


**Figure S15.** Heatmap of drug response result of live cell fluorescence intensity ($FI_{blue}$) from BhiCC framework-embedded spheroids from HepG2 (top) and LNCaP (bottom) after DOX treatment (24 h treatment and 24 h incubation). Each column represents DOX concentrations from 0 to 200 μM, respectively, while first to fourth rows represent experimental replicates (*i.e.*, Sample #1–#4; $N = 4$).

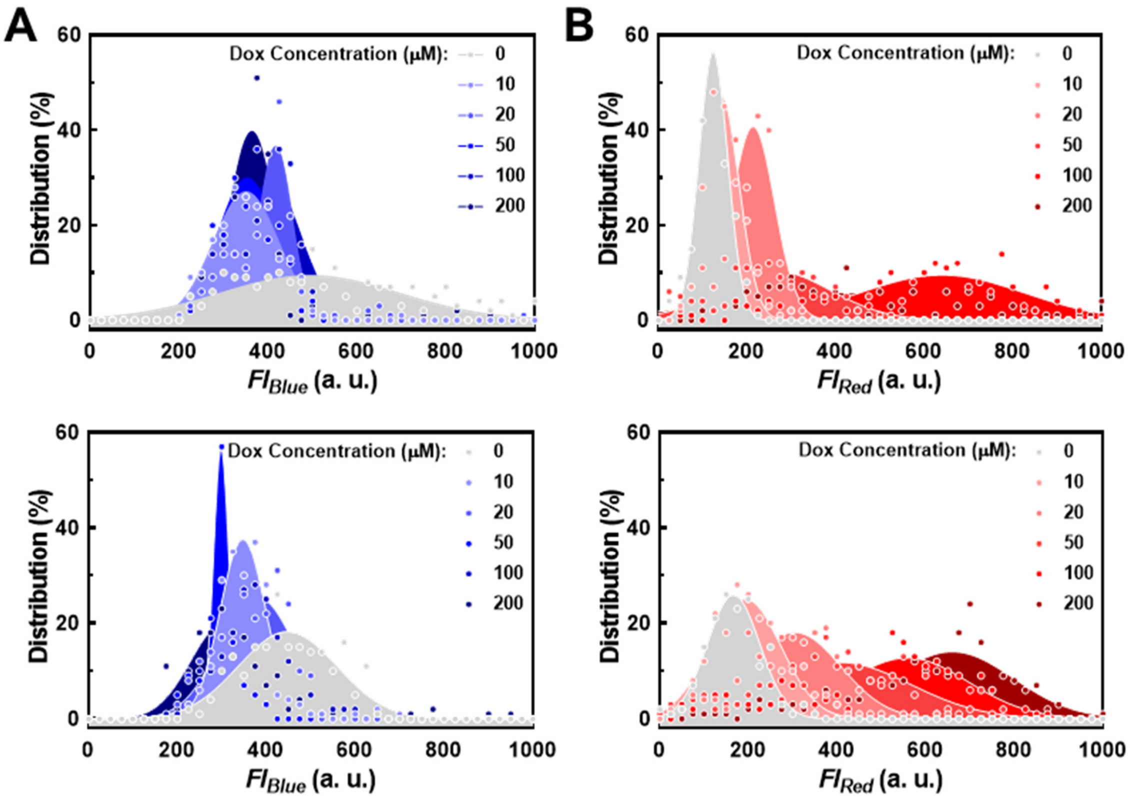


**Figure S16**. Histogram and Gaussian fitting results from drug response analysis of BhiCC framework-embedded spheroids derived from HepG2 (top of each panel) and LNCaP (bottom of each panel) cells, as measured by a plate reader. **(A)** The live cell signal ($FI_{blue}$) from the spheroids and **(B)** the dead cell signal ($FI_{red}$) resulting from drug response. Scattered dots represent the frequency distribution of each fluorescence intensity, while the filled areas indicate the respective fitted Gaussian plots.

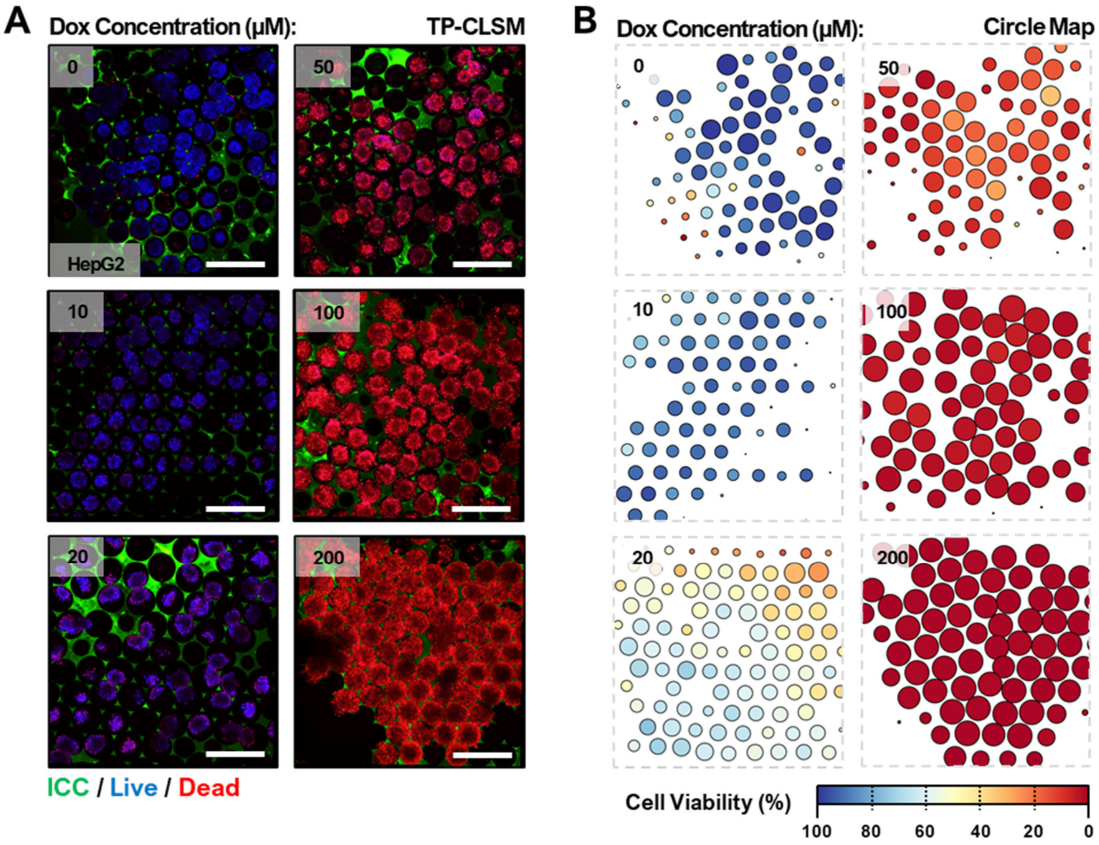


**Figure S17.** Proof-of-concept statistically reliable high-content (HC) drug screening with BhiCC framework. **(A)** Single *Z*-plane TP-CLSM images of HepG2 spheroids within BhiCC framework after DOX treatment (24 h treatment and 24 h incubation; green: 5-DTAF on F-LMA, blue: live cells, and red: dead cells; scale bar: 500 μm). **(B)** Simplified circle mapping images of HepG2 spheroids within BhiCC framework after DOX treatment (24 h treatment and 24 h incubation).

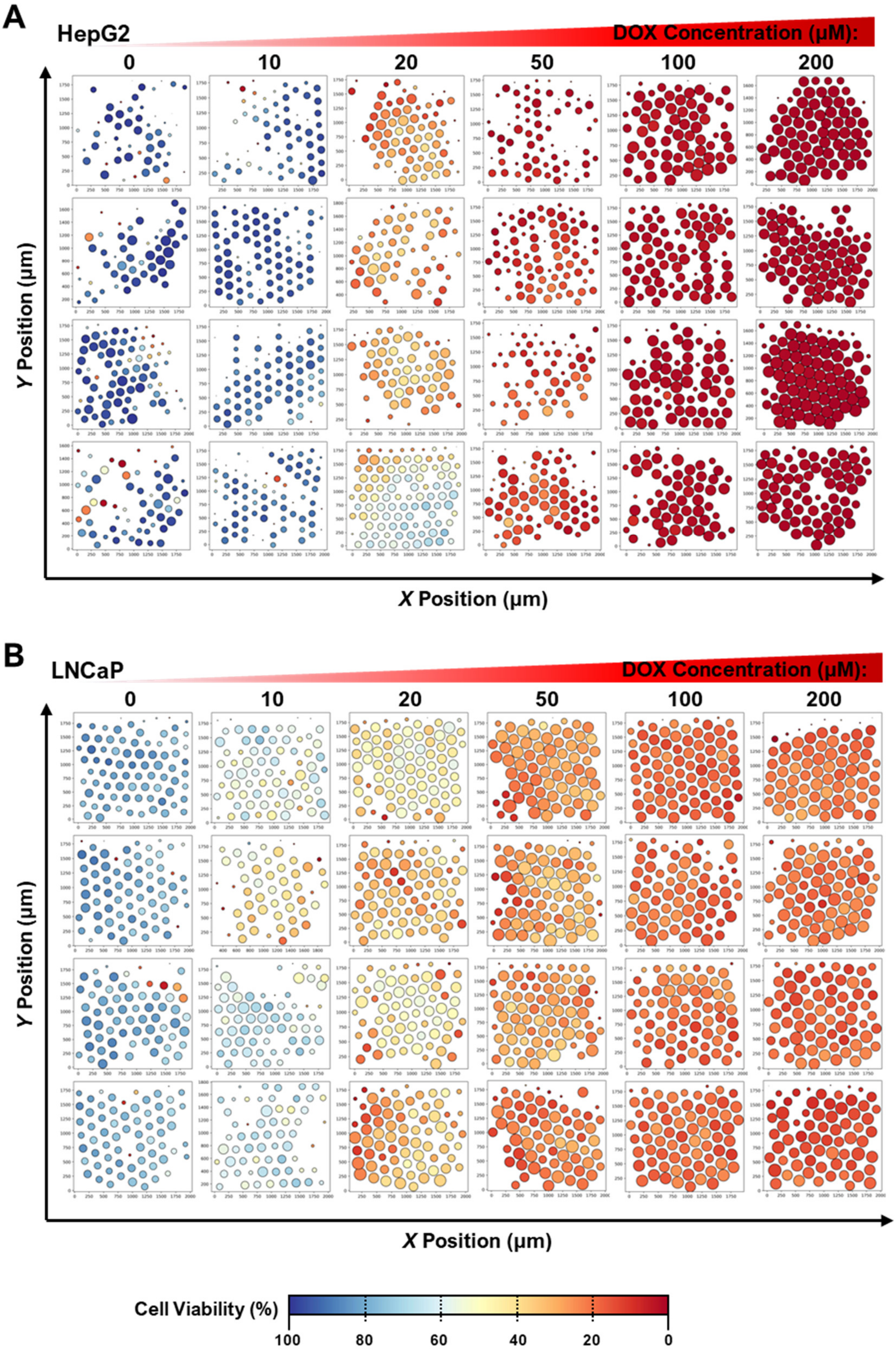


**Figure S18.** Full image set of simplified circle mapping images of **(A)** HepG2 and **(B)** LNCaP spheroids within BhiCC framework after DOX treatment (24 h treatment and 24 h incubation).

#### Supplementary Tables

**Table S1.** Statistical description for HC drug screening with BhiCC framework.

| Drug Response (%) | | 0 µM | 10 µM | 20 µM | 50 µM | 100 µM | 200 µM |
|---|---|---|---|---|---|---|---|
| **HepG2** | # of Spheroids (ea) | **251** | **316** | **217** | **288** | **285** | **320** |
| | Mean | 12.71 | 27.07 | 70.38 | 92.895 | 98.786 | 99.3506 |
| | Std. Deviation (SD) | 14.63 | 11.39 | 6.926 | 1.774 | 0.2737 | 0.1473 |
| | Std. Error of Mean (SEM) | 0.9237 | 0.6406 | 0.4702 | 0.1045 | 0.0162 | 0.008 |
| **LNCaP** | # of Spheroids (ea) | **298** | **267** | **315** | **317** | **312** | **374** |
| | Mean | 18.53 | 36.19 | 43.22 | 60.5 | 67.22 | 73.79 |
| | Std. Deviation (SD) | 3.968 | 4.277 | 6.281 | 5.348 | 2.716 | 3.154 |
| | Std. Error of Mean (SEM) | 0.2299 | 0.2618 | 0.2527 | 0.3004 | 0.1537 | 0.1631 |

**Table S2.** The *P*-values and area under the curve (AUC) values from receiver operating characteristic (ROC) analysis for every data point collected from BhiCC framework-based HC drug screening. HepG2 (top) and LNCaP (bottom). ****: $P<0.0001$. Number for each plot represents AUC value from each comparison.

| HepG2 | versus | | DOX Concentration (µM) | | | | | |
|---|---|---|---|---|---|---|---|---|
| | | | 0 | 10 | 20 | 50 | 100 | 200 |
| DOX Concentration (µM) | 0 | *P* | - | **** | **** | **** | **** | **** |
| | | AUC | - | 0.7746 | 0.9454 | 0.9831 | 0.9916 | 0.9947 |
| | 10 | *P* | | - | **** | **** | **** | **** |
| | | AUC | | - | 0.9449 | 0.9919 | 0.9967 | 0.9968 |
| | 20 | *P* | | | - | **** | **** | **** |
| | | AUC | | | - | 0.9493 | 0.9962 | 0.9967 |
| | 50 | *P* | | | | - | **** | **** |
| | | AUC | | | | - | 0.8787 | 0.9748 |
| | 100 | *P* | | | | | - | **** |
| | | AUC | | | | | - | 0.9101 |
| | 200 | *P* | | | | | | - |
| | | AUC | | | | | | - |

| LNCaP | versus | | DOX Concentration (µM) | | | | | |
|---|---|---|---|---|---|---|---|---|
| | | | 0 | 10 | 20 | 50 | 100 | 200 |
| DOX Concentration (µM) | 0 | *P* | - | **** | **** | **** | **** | **** |
| | | AUC | - | 0.9541 | 0.9928 | 0.9952 | 0.9965 | 0.9966 |
| | 10 | *P* | | - | **** | **** | **** | **** |
| | | AUC | | - | 0.8782 | 0.9783 | 0.9899 | 0.9913 |
| | 20 | *P* | | | - | **** | **** | **** |
| | | AUC | | | - | 0.9075 | 0.9687 | 0.9687 |
| | 50 | *P* | | | | - | **** | **** |
| | | AUC | | | | - | 0.62 | 0.8149 |
| | 100 | *P* | | | | | - | **** |
| | | AUC | | | | | - | 0.5998 |
| | 200 | *P* | | | | | | - |
| | | AUC | | | | | | - |
